# Priors for Genotyping Polyploids

**DOI:** 10.1101/751784

**Authors:** David Gerard, Luís Felipe Ventorim Ferrão

## Abstract

**Motivation:** Empirical Bayes techniques to genotype polyploid organisms usually either (i) assume technical artifacts are known *a priori* or (ii) estimate technical artifacts simultaneously with the prior genotype distribution. Case (i) is unappealing as it places the onus on the researcher to estimate these artifacts, or to ensure that there are no systematic biases in the data. However, as we demonstrate with a few empirical examples, case (ii) makes choosing the class of prior genotype distributions extremely important. Choosing a class that is either too flexible or too restrictive results in poor genotyping performance.

**Results:** We propose two classes of prior genotype distributions that are of intermediate levels of flexibility: the class of proportional normal distributions and the class of unimodal distributions. We provide a complete characterization of and optimization details for the class of unimodal distributions. We demonstrate, using both simulated and real data, that using these classes results in superior genotyping performance.

**Availability and Implementation:** Genotyping methods that use these priors are implemented in the updog R package available on the Comprehensive R Archive Network: https://cran.r-project.org/package=updog. All code needed to reproduce the results of this paper is available on GitHub: https://github.com/dcgerard/reproduce_prior_sims.

## 1 Introduction

Because of their importance in agriculture [Udall and Wendel, 2006], evolution [Soltis et al., 2014], and biodiversity [Soltis and Soltis, 2000], scientists are increasingly interested in studying the genomic landscape of polyploid organisms (organisms with more than two copies of their genome). In these studies, researchers usually first estimate the number of copies of each allele (dosage) at given polymorphic loci in a sample. These dosages may then be fed into downstream analyses such as genome-wide association studies [Rosyara et al., 2016, Ferrão et al., 2018, Benevenuto et al., 2019], genomic prediction [Endelman et al., 2018, Amadeu et al., 2019, de Bem Oliveira et al., 2019, Lara et al., 2019], and genetic mapping [Shirasawa et al., 2017, Ferreira et al., 2019]. To determine allele dosage, researchers first measure the reference and alternative counts of small reads from some sequencing technology [Baird et al., 2008, Elshire et al., 2011]. Based on these read counts, they then infer the dosage of reference and alternative alleles using statistical methods.

Due to constraints in sequencing depth, many genotyping methods incorporate empirical Bayes techniques to borrow strength between individuals [Voorrips et al., 2011, Serang et al., 2012, Maruki and Lynch, 2017, Blischak et al., 2018, Gerard et al., 2018, Clark et al., 2019]. These methods posit some class of prior distributions for the possible genotypes, select a distribution among this class using maximum marginal likelihood, and incorporate this prior genotype distribution in a Bayesian inference scheme to derive a posterior genotype distribution from which genotype calls are made. Adaptively estimating the prior distribution allows methods to automatically gauge the level of prior information in the data and use this prior information to improve genotyping.

Besides the genotype of the individual, there are usually other, more technical, aspects of the data that should be taken into consideration during allele dosage inference, including sequencing error rate, allele bias, and overdispersion [Gerard et al., 2018]. A principled empirical Bayes approach would integrate over the uncertainty in these other parameters prior to estimating the genotype prior. However, computation is a major concern as there are often tens or hundreds of thousands of SNPs to genotype in a single dataset, and even increases of seconds can make a genotyping method untenable for applied researchers. Thus, for computational reasons, genotyping methods either assume these parameters are known *a priori* [Maruki and Lynch, 2017, Blischak et al., 2018, Clark et al., 2019] or estimate them simultaneously with the prior genotype distribution [Voorrips et al., 2011, Gerard et al., 2018].

Assuming these parameters are known *a priori* is unappealing as it prohibits adapting to the features present in the data. However, using the second strategy of simultaneously estimating technical features with the genotype prior makes the choice of the class of priors of vital importance. To demonstrate this, we present a genotype plot [Gerard et al., 2018] of a single SNP from the autotetraploid potato data from Uitdewilligen et al. [2013] in panel A of Figure 1. A genotype plot contains the reference counts of an individual on the *y*-axis and the alternative counts on the *x*-axis. Individuals with a dosage of four reference alleles would lie near the *x* = 0 vertical line, while individuals with a dosage of zero reference alleles would lie near the *y* = 0 horizontal line. The plot seems to display two clusters of individuals, which would indicate to a researcher that there is a group of potatoes with a genotype of four reference alleles and a group of potatoes with a genotype of three reference alleles. However, when we fit updog (the software implementation of the empirical Bayes genotyping method described in Gerard et al. [2018]) to these data, we obtain unreasonable genotype estimates when we either allow the class of priors to be anything over genotypes {0, 1, 2, 3, 4} (panel B of Figure 1) or we constrain the prior to be the discrete uniform distribution over genotypes {0, 1, 2, 3, 4} (panel C of Figure 1). These results generalize to other datasets (Supplementary Figure S1). (As a preview, panel D of Figure 1 demonstrates that using one of our proposed classes of prior distributions results in intuitive genotype estimates).

**Figure 1:**
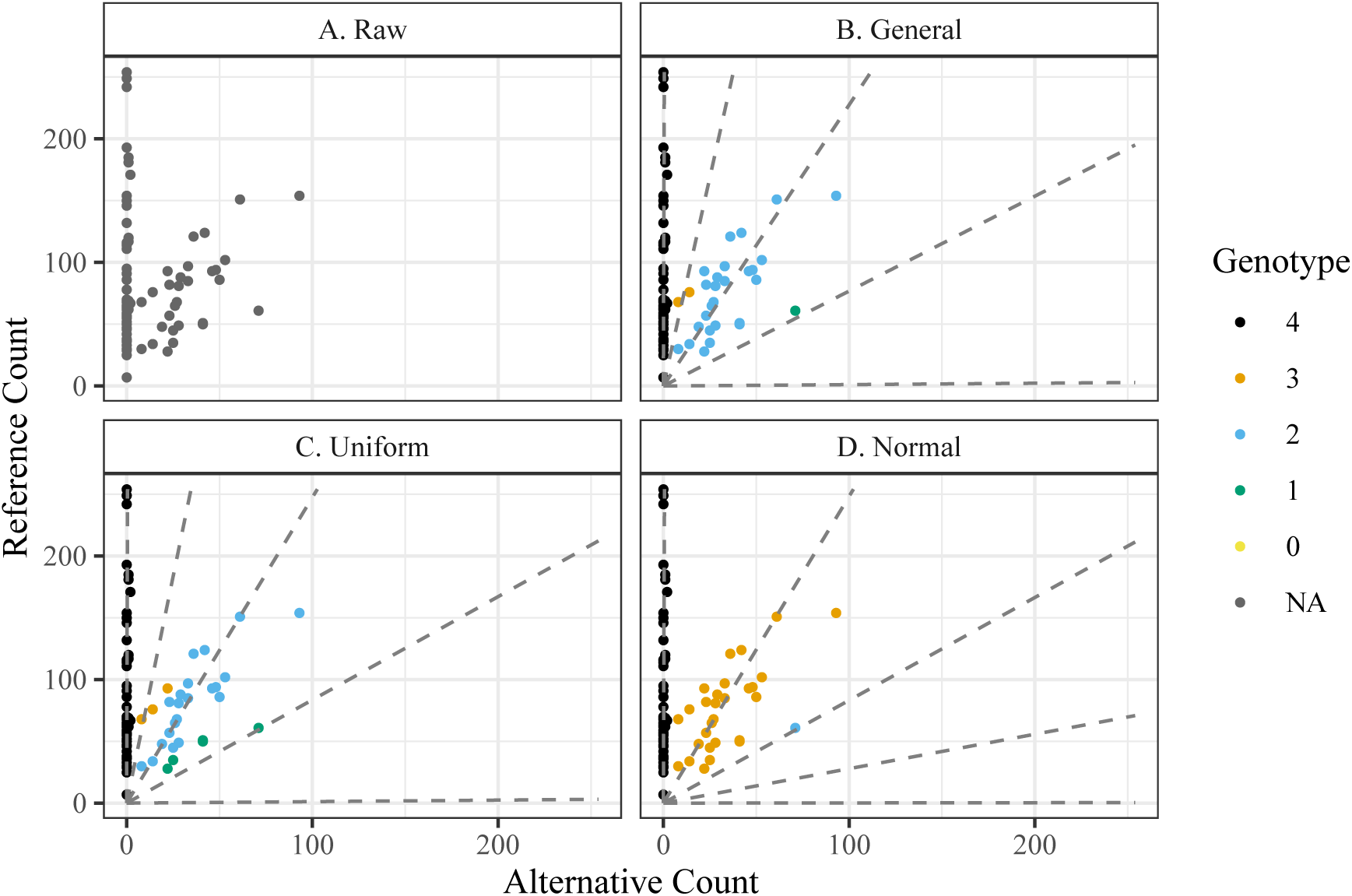
The number of reference reads (*y*-axis) versus the number of alternative reads (*x*-axis) from 84 autotetraploid potatoes for a SNP from Uitdewilligen et al. [2013]. The raw unannotated data are presented in panel (A). The other panels are color-coded by estimated genotype according to updog fits using either (B) the general class of priors, (C) the discrete uniform prior distribution, or (D) the class of proportional normal priors. The dashed lines in panels (B) through (D) are the estimated expected counts, and are functions of the estimated allele bias and estimated sequencing error rate [see Gerard et al., 2018]. Only the proportional normal class of priors provides intuitive genotyping.

This indicates that using a class of priors that is either too flexible or too restrictive can result in poor genotyping. In the case of the discrete uniform prior, updog effectively tries to estimate the allele bias and sequencing error to make the empirical genotype distribution as close to a discrete uniform as possible. As the real genotype distribution is not discrete uniform, this results in extreme estimates of the allele bias and unreasonable genotype estimates. In the case of the general prior, since we do not have a data-adaptive prior distribution on the bias, there is little loss in marginal likelihood for having an extreme allele bias. And so the bias parameter is free to vary as long as it increases the marginal likelihood. Indeed, if we include a slight penalty on the bias parameter, the general class of prior distributions results in reasonable genotype estimates (Supplementary Figure S2). These results indicate that having too flexible a class of prior distributions can result in unstable genotype estimates; while having too constrained a class of prior distributions can result in the likelihood parameters adapting to the prior distribution instead of having the prior distribution improve our estimates of the genotypes.

In this paper, we propose two classes of prior distributions for genotyping polyploids which are designed to have intermediate levels of flexibility (Section 2). One is the class of discrete unimodal distributions, for which we provide a complete characterization. The other is the class of distributions whose probability masses are proportional to a normal density. We provide optimization details for these classes, along with other classes of priors used in the literature. We demonstrate, through simulation and on real data, that these two classes, and in particular the proportional normal class, is the most robust to varying genotype distributions (Section 3). The proportional normal prior is the default in the updog software available on the Comprehensive R Archive Network: https://cran.r-project.org/package=updog.

## 2 Methods

We now provide a brief description of the model and genotyping procedure we study. We assume that all inference is done one SNP at a time (though see the discussion in Section 4). Let *x*_*i*_ be the reference read-counts for individual *i*, let *n*_*i*_ be the total number of reads for individual *i*, let *K* be the ploidy of the species under study, let *y*_*i*_ ∈ {0, 1, …, *K*} be the (unknown) genotype for individual *i*, let ***η*** be a vector of likelihood-specific parameters (such as the allele bias, sequencing error rate, or overdispersion), and let ***θ*** be a vector of prior-specific parameters. Different values of ***θ*** index the class of allowable genotype distributions. Let *f* (*x*_*i*_|*y*_*i*_, *n*_*i*_, ***η***) denote the likelihood of *x*_*i*_. This could, for example, be the likelihood from Voorrips et al. [2011] or Gerard et al. [2018], but we allow it to be more general. Let *f* (*y*_*i*_|***θ***) denote the prior probability of genotype *y*_*i*_. We suppose that the inference strategy consists of the following two steps: First, estimate the likelihood- and prior-specific parameters by maximum marginal likelihood:

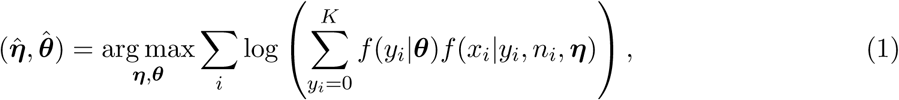

and second, derive the posterior distribution of the genotypes for each individual:

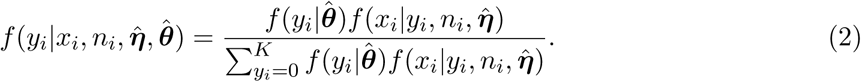

These posterior probabilities are then used to make genotype calls. Deriving a general-purpose expectation-maximization algorithm [Dempster et al., 1977] to obtain 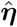 and 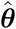 is not difficult (Supplementary Section S2.1), though special optimization considerations are needed for some of the priors we discuss in this section (Supplementary Sections S2.1 and S2.2).

Prior distributions that have been used in the past (though not necessarily in the above framework) include:

1. The discrete uniform distribution [McKenna et al., 2010]:

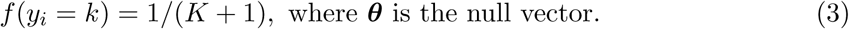
2. The distribution of autopolyploid sibling genotypes [Serang et al., 2012]:

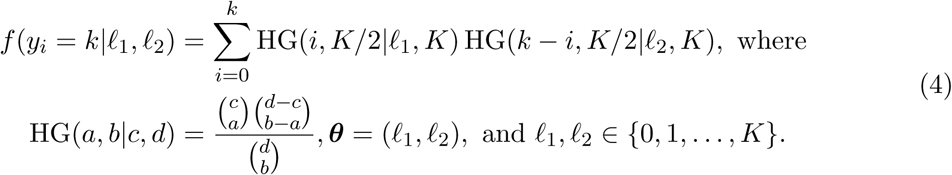 Equation (4) is merely a convolution of two hypergeometric distributions. The *ℓ*_1_ and *ℓ*_2_ parameters are the genotypes of the two parents. We will denote this distribution as the “F1 distribution” as it results from an F1 cross of autopolyploid individuals.
3. The binomial distribution [Martin et al., 2010, Li, 2011]:

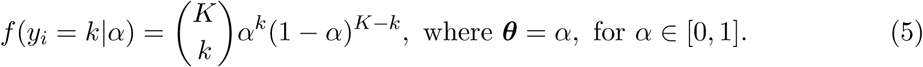 The *α* parameter is the allele frequency of the reference allele. The binomial distribution results by assuming the individuals are in Hardy-Weinberg equilibrium.
4. The beta-binomial distribution [Blischak et al., 2018]:

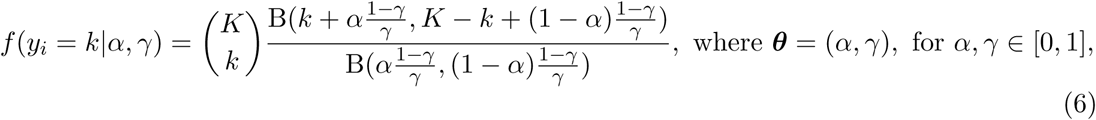

where B(*a, b*) represents the beta function with parameters *a* and *b*. The *α* parameter is the allele frequency of the reference allele, while *γ* is an overdispersion parameter. This is slightly different than the prior used in Blischak et al. [2018] as there the overdispersion parameter is individual-specific, while in (6) it is locus-specific.
5. The general categorical distribution [DePristo et al., 2011, Li, 2011]:

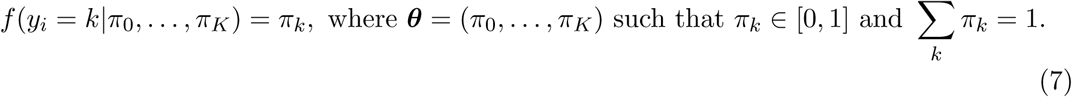

There are many issues with these previously used priors. The issues with the discrete uniform and general distributions were already noted in Section 1. The F1 and binomial distributions are only applicable in specific situations. The F1 distribution can only be applied if the individuals are known siblings, the binomial distribution relies on the strong assumptions of Hardy-Weinberg equilibrium [Crow, 1988]. Though the beta-binomial distribution can account for overdispersion, it is unable to account for *under* dispersion, which may be caused by, e.g., unobserved relatedness (for example, the F1 distribution is generally less dispersed than a binomial, and hence a beta-binomial).

In order to account for both over- and underdispersion, in our first proposal we model genotype probabilities as being proportional to a normal density:

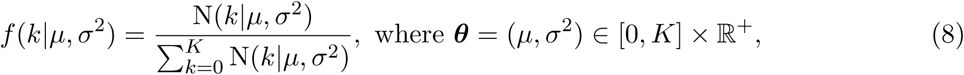

where N(⋅|*µ, σ*^2^) represents the normal density with mean *µ* and variance *σ*^2^. This distribution, though somewhat ad hoc, can approximate very well distributions from the classes of binomial, beta-binomial, and F1 distributions (Supplementary Figure S3). One can thus essentially consider it a general class that encompasses as special cases these classes of distributions. Varying *σ*^2^ allows for the prior distribution to accommodate both over- and underdispersion. We call this the “proportional normal distribution”.

Motivated by the fact that most of the prior distributions previously proposed in the literature are unimodal, our second proposal is to model genotype probabilities within the flexible class of unimodal probability mass functions. In Supplementary Section S1 we exactly characterize this class of distributions using mixtures of discrete uniform distributions. This idea was motivated by the work of Stephens [2016] who approximated continuous densities using mixtures of continuous uniform distributions. Our characterization lends itself to a convex optimization problem during the M-step of the EM algorithm (Supplementary Section S2.1). But rather than use standard convex optimization techniques, we found it faster to develop a weighted EM scheme to optimize over the class of unimodal distributions (Supplementary Section S2.2).

All priors explored in this paper are summarized in Supplementary Table S2.

## 3 Results

### 3.1 Simulation results

In this section we compare, through simulation, the seven different classes of prior distributions from Section 2. Our strategy was to generate genotypes under seven different genotype distributions, where each genotype distribution was designed to be favorable to one of the classes of prior distributions. The discrete uniform, binomial (Hardy-Weinberg), beta-binomial, and F1 favorable genotype distributions were all generated according to their assumed priors. For the genotype distribution favorable to the proportional normal prior, we generated genotypes according to the proportional normal distribution with a small variance since we motivated this distribution by the need to account for underdispersion. For the unimodal favorable genotype distribution, we allowed genotype probabilities to decline linearly from *K* to 0 since none of the other more structured prior classes have such linear declines. Finally, for the genotype distribution favorable to the general prior, we specified non-unimodal genotype proportions, which none of the other priors should be able to accommodate. The specific genotype distributions are listed in Supplementary Section S3 and are graphically represented in Supplementary Figure S6.

Given individual genotypes, we generated sequencing data according to the likelihood in Gerard et al. [2018], setting the sequencing error rate to 0.001, varying the overdispersion parameter over {0, 0.005, 0.01} (where larger values indicate greater dispersion), and varying the allele bias parameter over {1, 0.75, 0.5} (where values further from 1 indicate greater bias). Each repetition, we generated thirty hexaploid individuals at a sequencing depth of 100 reads. For each unique combination of allele bias, overdispersion, and genotype distribution, we generated 500 datasets. For each dataset, we fit updog assuming one of the seven prior classes described in Section 2.

The results under conditions of no allele bias and no overdispersion are presented in Figures 2 and 3. The results under varying levels of bias and overdispersion are presented in Supplementary Figures S7–S27. Figure 2 plots the proportion of individuals genotyped correctly (with larger values being better) while Figure 3 plots the difference between the estimated proportion of individuals misclassified and the actual proportion of individuals misclassified (with values closer to 0 being better). In Figures 2 and 3, we see that each class of prior distributions performs best when the genotype distribution is designed to be favorable to it. Except for the classes of proportional normal distributions and unimodal distributions, every class performs very poorly under some genotype distribution. However, the class of unimodal distributions performs poorly under some genotype distributions under larger levels of bias and overdispersion (Supplementary Figures S8, S11, S15, and S18). The class of proportional normal distributions only performs poorly under the (unrealistic) general-favorable genotype distribution with large levels of bias and overdispersion (Supplementary Figures S13 and S20). Only the class of proportional normal priors performs adequately when the gentoypes are very underdispersed (Supplementary Figures S11 and S18). Estimates of the allele bias when using the class of proportional normal priors appear to be unbiased (Supplementary Figures S23–S27). These results indicate that the class of proportional normal distributions performs the best under a wide variety of settings.

**Figure 2:**
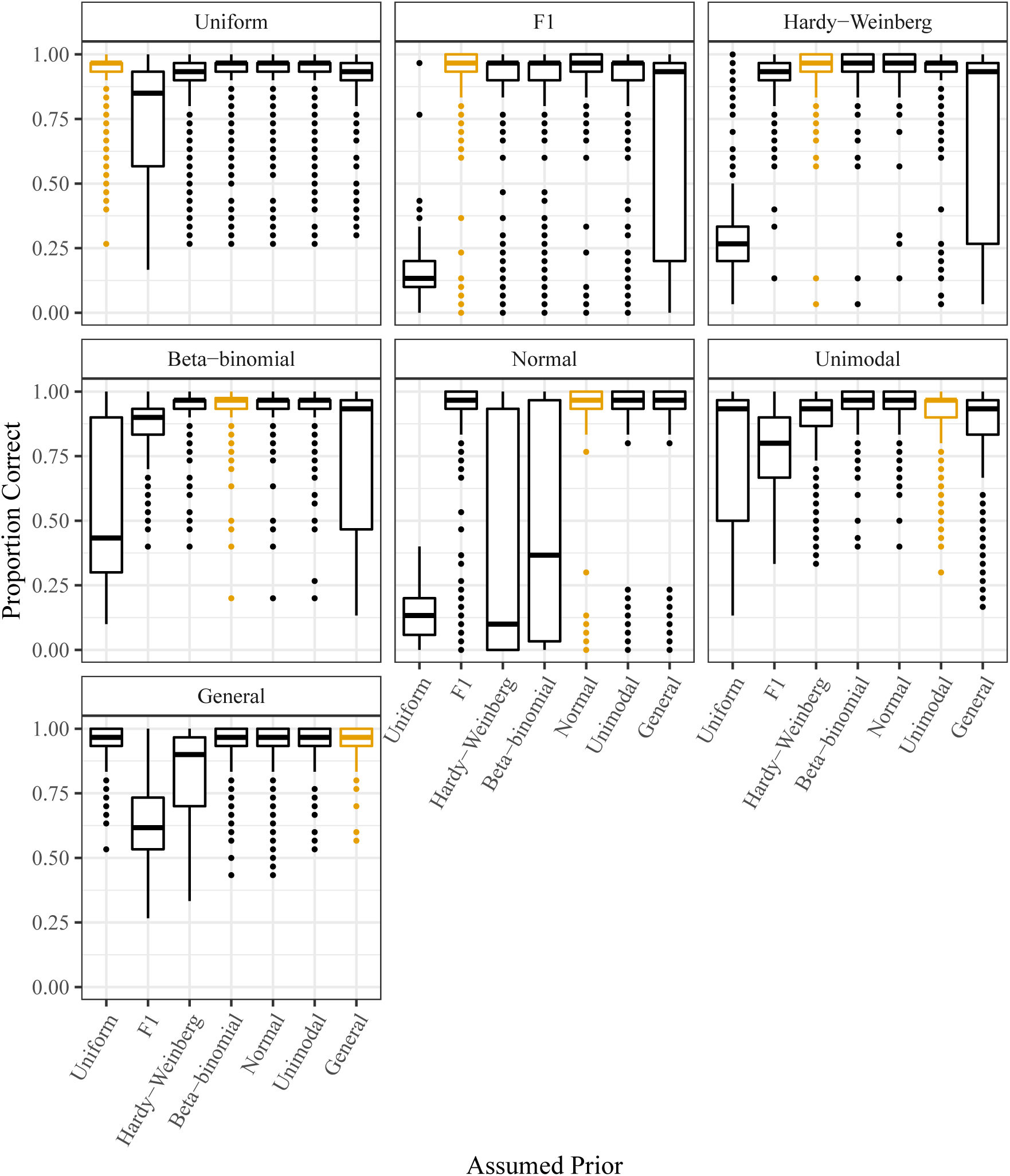
Proportion of individuals genotyped correctly (*y*-axis) stratified by the assumed class of prior distributions (*x*-axis) used in an updog fit. The facets index the distributions used to generate the genotypes, with each genotype distribution designed to be favorable to one of the classes of prior distributions (orange). The distributions are ordered roughly from least flexible (the discrete uniform) to most flexible (the general class). The read counts were generated assuming no allele bias and no overdispersion.

**Figure 3:**
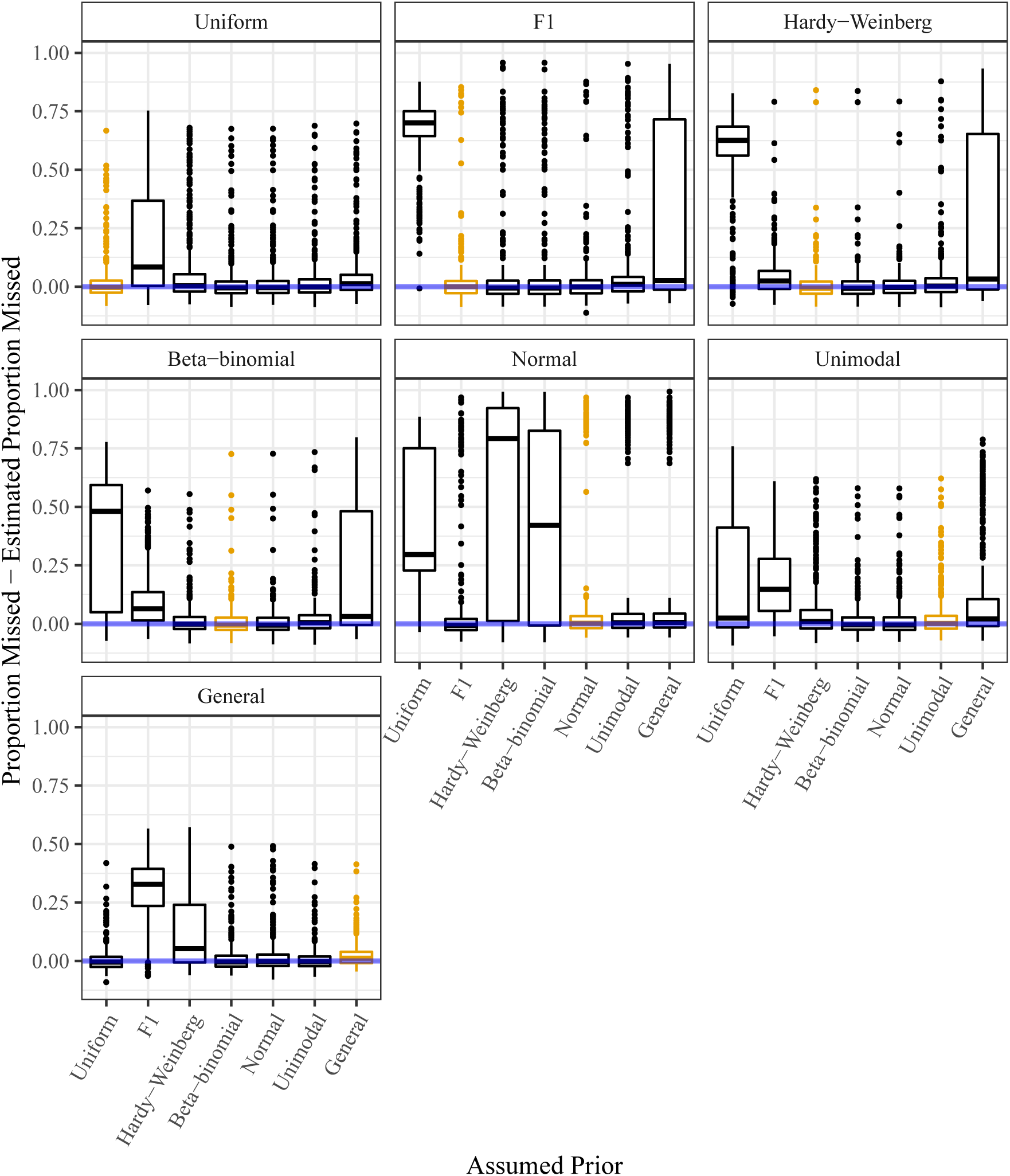
Estimated proportion of individuals incorrectly genotyped subtracted from the actual proportion of individuals incorrectly genotyped (*y*-axis) stratified by the assumed class of prior distributions (*x*-axis) used in an updog fit. The horizontal line (at *y* = 0) indicates unbiased estimation of the misclassification error rate. The facets index the distributions used to generate the genotypes, with each genotype distribution designed to be favorable to one of the classes of prior distributions (orange). The distributions are ordered roughly from least flexible (the discrete uniform) to most flexible (general). The read counts were generated assuming no allele bias and no overdispersion.

### 3.2 Sweet potato results

In this section, we compare the seven classes of prior distributions discussed in Section 2 using the data from Shirasawa et al. [2017]. The hexaploid sweet potatoes in these data are known siblings, resulting from a single generation of selfing (an S1 cross). Thus, the class of genotype distributions is known to be F1 (4) with the constraint that *ℓ*_1_ = *ℓ*_2_ [Gerard et al., 2018], which we will simply call “S1”. We fit updog, using all seven prior classes in Section 2 as well as the S1 prior class, to the 1000 SNPs with mean read-depth closest to 100. We ran updog only on a sample of twenty individuals, as having fewer individuals makes the class of prior distributions most important. For each SNP, we calculated the Euclidean distances between all pairs of posterior mean genotypes resulting from the eight different fitted prior classes. We also calculated the Hamming distances (divided by the number of individuals) between the posterior mode genotypes resulting from the eight different fitted prior classes.

The results are presented in Figure 4. As the genotype distribution is known to belong to the S1 class, we will consider its genotypes a proxy for the true genotypes. Thus, prior classes that are closest to S1 (in terms of either Euclidean distance from posterior mean genotypes, or Hamming distance from posterior mode genotypes) perform the best. We see in Figure 4 that F1 performs the best. This is expected since S1 is a small subclass of the class of F1 distributions. The second best performance comes from the class of proportional normal priors. Thus, on real data, we see the gains attained by using the class of proportional normal priors. Mean distances between the results of all methods are presented in Supplementary Figures S28 and S29.

**Figure 4:**
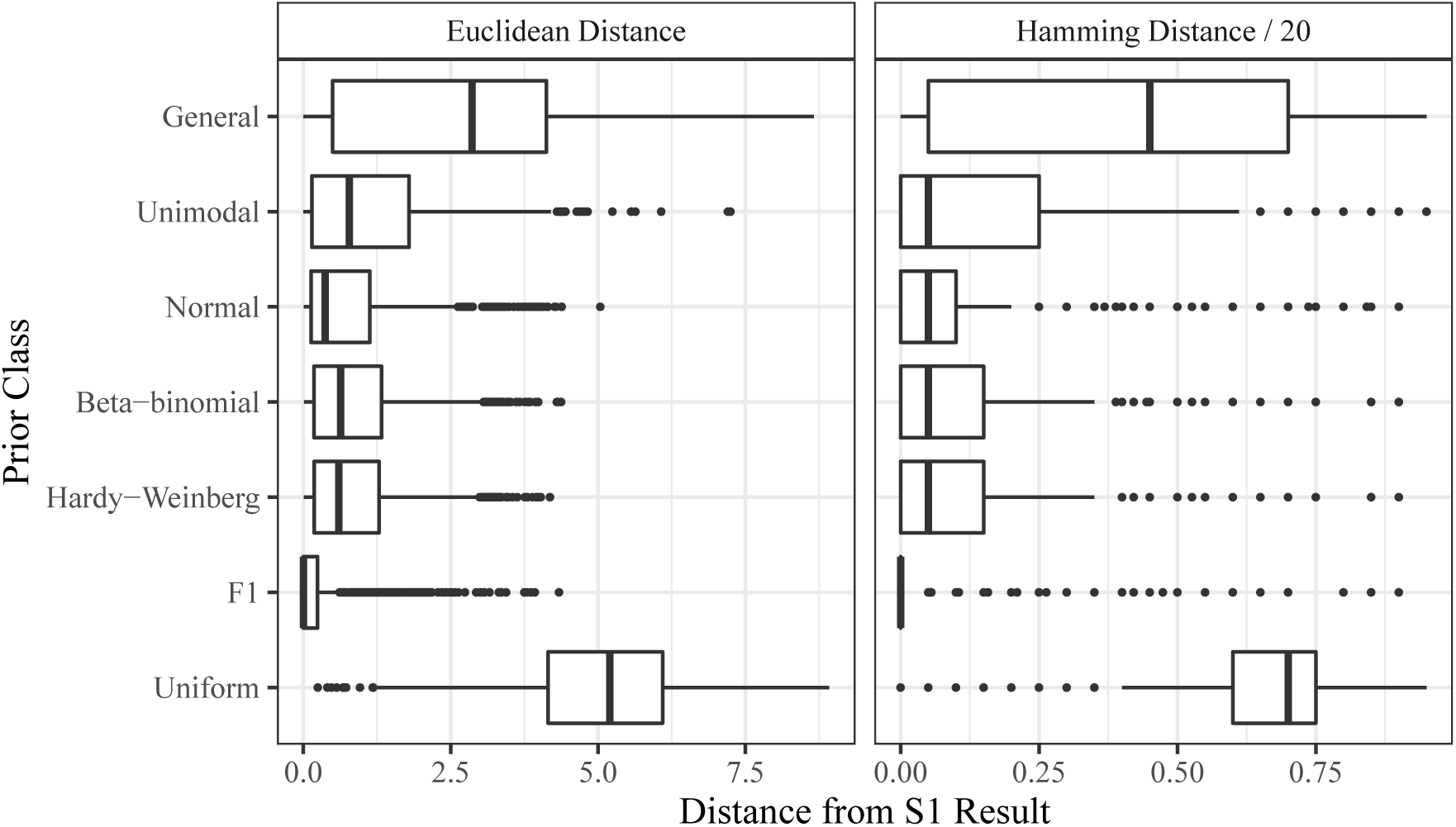
Boxplots of distances (*x*-axis) stratified by prior class (*y*-axis). The left facet contains the Euclidean distances between posterior means when either using the S1 prior or the specified prior on the *y*-axis. The right facet contains the proportion of individuals genotyped differently when either using the S1 prior or the specified prior on the *y*-axis. Distances closer to 0 indicate superior performance.

## 4 Discussion

In this paper, we reviewed the classes of prior distributions used to genotype polyploids, found these classes either too flexible or too restrictive, and proposed two new classes of priors of intermediate levels of flexibility. This required us to characterize the class of discrete unimodal distributions using mixtures of discrete uniform distributions. We then demonstrated, through simulation, that our proposals work better under the genotyping inference scheme in updog. In particular, we recommend to use by default the class of proportional normal distributions as it resulted in robust performance under different gentoype distributions. The proportional normal prior is the default in the updog software available on the Comprehensive R Archive Network: https://cran.r-project.org/package=updog. Our classes of priors are also more generally applicable to other inference schemes.

Our priors are appropriate defaults for a wide variety of genotyping scenarios. For example, breeding populations might exhibit violations in Mendelian segregation proportions (4) due to double reduction [Stift et al., 2010], preferential pairing [Voorrips and Maliepaard, 2012], or resulting from an allele being semi-lethal. Additionally, unstructured populations might exhibit violations from Hardy-Weinberg equilibrium (5) due to selection or non-random mating [Crow, 1988]. Our proportional normal and unimodal classes of priors may be used in all of these scenarios without explicitly modeling these processes.

Here, we only analyzed one SNP at a time, though some recent approaches have also attempted to borrow strength between SNPs [Blischak et al., 2018, Clark et al., 2019]. In particular, Clark et al. [2019] tries to account for linkage disequilibrium (LD) between SNPs. However, their method, though intuitive, is somewhat ad hoc. Generalizing our proportional normal prior would allow for a more principled approach to accounting for LD by simply letting the genotype distribution across multiple SNPs be proportional to a multivariate normal. To account for the high-dimensionality present due to the large number of SNPs, one could impose strong structural constraints on the covariance matrix, such as letting it decay exponentially according to recombination rate [Wen and Stephens, 2010].

## Data availability

All methods are implemented in the `updog` software available on the Comprehensive R Archive Network: https://cran.r-project.org/package=updog. All code and instructions to reproduce the results of this paper are available on GitHub: https://github.com/dcgerard/reproduce_prior_sims.

## Acknowledgments

All graphics were made using ggplot2 [Wickham, 2016] in the R statistical language [R Core Team, 2019].

We thank the College of Arts and Sciences at American University for providing a travel grant from the Mellon Fund. This allowed the authors to meet and collaborate on this manuscript.

## S1 Characterizing the class of discrete unimodal distributions

### Theorem 1.

*Let* 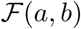 *denote the set of integers between a and b. That is*,

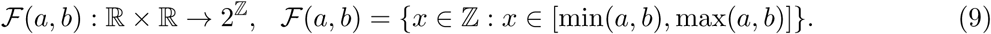

*Then a probability mass function f* (⋅) *with support on* {0, 1, …, *K*} *is unimodal if and only if there exists an a* ∈ (−1, *K* + 1) \ {0, 1, …, *K*} *and* {*π*_0_, *π*_1_, …, *π*_*K*_} *with π*_*i*_ ∈ [0, 1] *and* 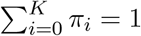 *such that*

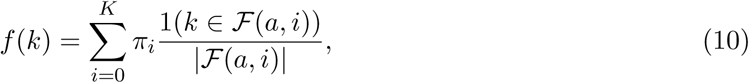

*where* 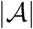 *denotes the number of elements in a set* 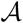 *and* 1(⋅) *is the indicator function*.

*Proof.* We will start by proving the “if” part. Suppose (10) holds. We must then prove three things: (i) *f* (⋅) is a probability mass function, (ii) if *k*_1_ *< k*_2_ *< a* then *f* (*k*_1_) ≤ *f* (*k*_2_), and (iii) if *k*_1_ *> k*_2_ *> a* then *f* (*k*_1_) ≤ *f* (*k*_2_). Note that *f* (*k*) is non-negative for all *k* since the summands in (10) are always non-negative. We also have

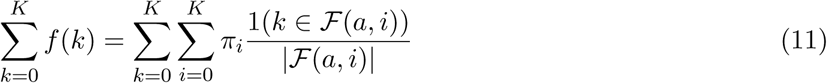

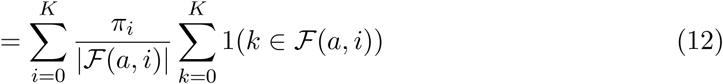

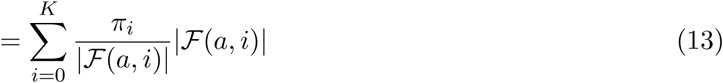

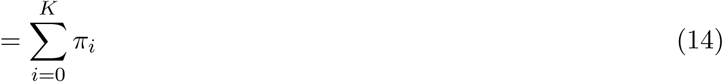

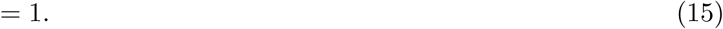

Thus, *f* (⋅) is a valid probability mass function. Now let *k*_1_ < *k*_2_ < *a*. Then

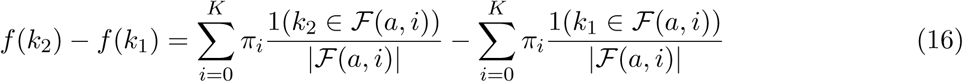

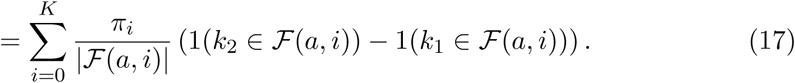

Note that if 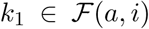 then 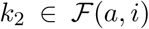, so 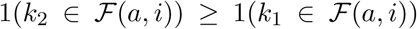. Thus, the summands in (17) are all non-negative, which implies that *f* (*k*_2_) ≥ *f* (*k*_1_). The proof for when *k*_1_ > *k*_2_ > *a* is similar. We have thus proved the “if” part.

We now prove the “only if” part. Suppose *f* (⋅) is a unimodal probability mass function with support on {0, 1*, …, K*}. Then there exists an *a* ∈ (−1*, K* + 1) \ {0, 1*, …, K*} such that *k*_1_ *< k*_2_ *< a* implies *f* (*k*_1_) ≤ *f* (*k*_2_) and *k*_1_ > *k*_2_ > *a* implies that *f* (*k*_1_) ≤ *f* (*k*_2_). For example, let *j* ∈ {0, 1, …, *K*} be any mode of *f* (⋅), then choose *a* = *j* + *ϵ* for any *ϵ* < 1/(*K* + 1). Now set

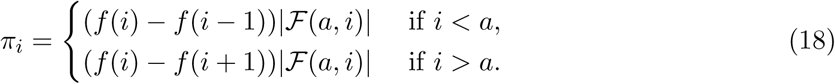

We must now show three things: (i) the definition of the *π*_*i*_’s used in (18) results in equality (10), (ii) the *π*_*i*_’s are all between 0 and 1, and (iii) the *π*_*i*_’s sum to 1. First, suppose that *k* < *a*. Then

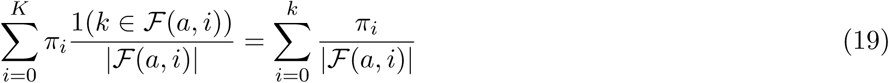

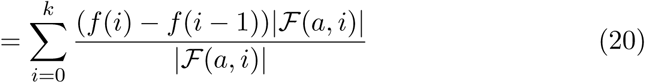

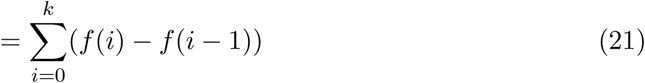

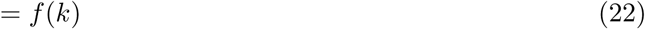

Equation (19) follows since 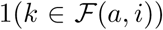 is 0 for *i* > *k* and is 1 for *i* ≤ *k*. Equation (22) follows from (21) by a telescoping argument and noting that *f* (−1) = 0. The argument for when *k* > *a* is similar. Thus, we have that (10) holds. By the unimodality of *f* (⋅), the *π*_*i*_’s in (18) are non-negative. Thus, it suffices to show that they sum to one. We have

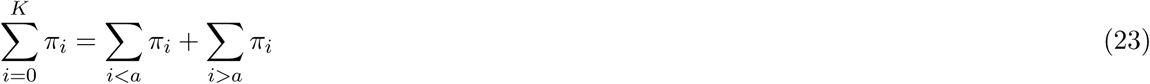

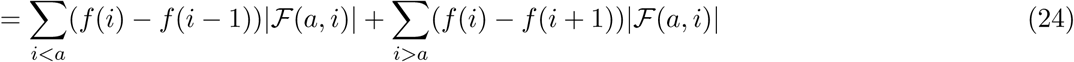

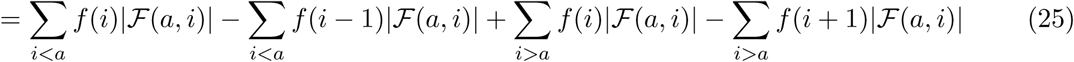

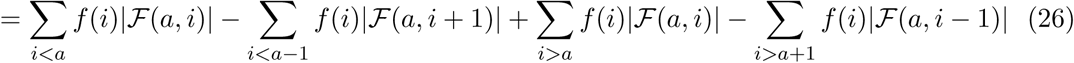

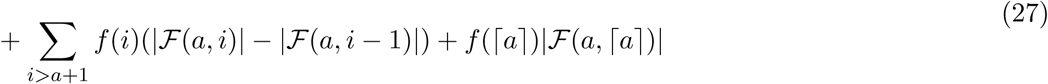

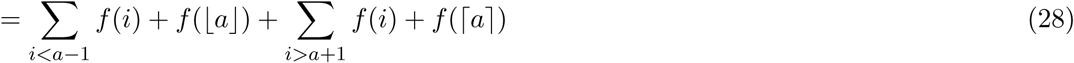

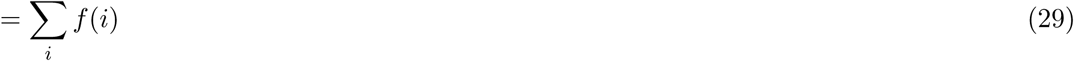

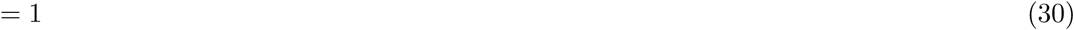

Equation (28) follows from the fact that 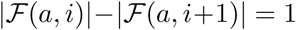 for *i* < *a*, 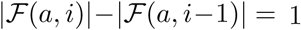 for *i* > *a*, and 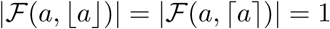.

## S2 Optimization details

### S2.1 EM derivation

We will now derive a general-purpose EM algorithm to solve (1). Let ***z***_*i*_ be a 1-of-(*K* + 1) vector indicating the genotype for individual *i*. Then the Complete log-likelihood of the ***z***_*i*_’s and *x*_*i*_’s is:

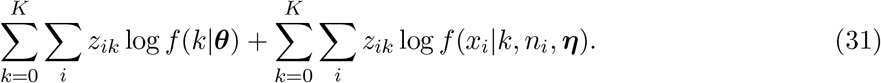

Using standard calculations, the posterior expectation of *z*_*ik*_ is

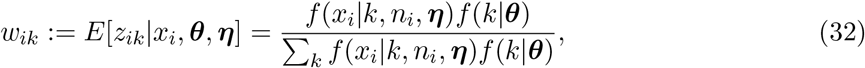

And the expected complete log-likelihood is

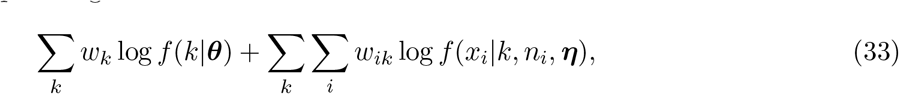

where *w*_*k*_:= Σ_*i*_*w*_*ik*_. Thus, the E-step of the EM algorithm consists of calculating the posterior expectations from (32). The M-step consists of two independent optimization problems, one for the likelihood-specific parameters and one for the prior-specific parameters:

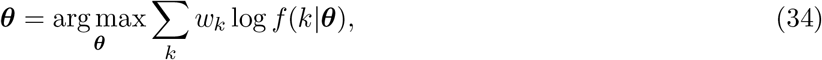

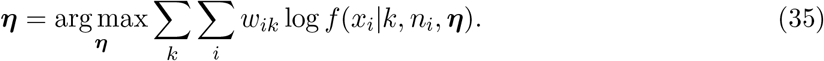

Since the E-step and the optimization in (35) is the same no matter the choice of class of priors, we focus on the optimization in (34).

We will now derive the optimization updates for the each of the classes of prior distributions discussed in Section 2. There is no prior optimization step for the discrete uniform distribution (3). For the beta-binomial (6) and proportional normal (8) distributions, we merely use gradient ascent to solve (34). For the binomial distribution (5), there exists a closed-form solution for update (34):

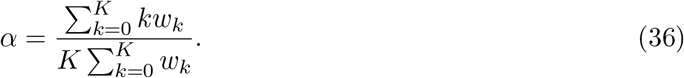

We also have a closed-form solution for the update with the general prior (7):

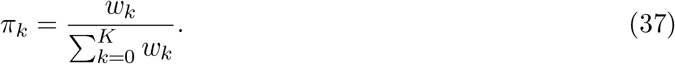

Note that Σ_*k*_ *w*_*k*_ in (36) and (37) is merely the number of individuals in the sample.

A naive update for the F1 distribution (4) would simply iterate over the possible levels of (*ℓ*_1_, *ℓ*_2_) ∈ {0, 1, …, *K*} × {0, 1, …, *K*}. However, this approach causes the resulting EM-algorithm to get stuck at certain levels of (*ℓ*_1_, *ℓ*_2_). To see this, recall that the prior optimization problem during the M-step is

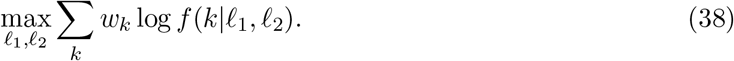

If *w*_*i*_ is non-zero during the current M-step, then moving to an (*ℓ*_1_, *ℓ*_2_) such that *f* (*i*|*ℓ*_1_, *ℓ*_2_) = 0 would result in a likelihood of −∞. Also, if the current values of (*ℓ*_1_, *ℓ*_2_) result in a non-zero *f* (*i*|*ℓ*_1_, *ℓ*_2_), then (32) implies that *w*_*i*_ can never become 0 during the E-step. This is major issue for the EM algorithm since the F1 distribution contains many exact zero probabilities, effectively inhibiting any exploration of (*ℓ*_1_, *ℓ*_2_). To remedy this, we introduce a small discrete uniform mixing component:

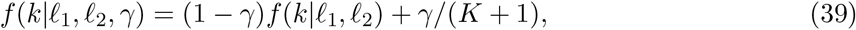

for some *γ* ∈ (0, 1). A principled approach would be to take *γ* to 0 during the EM-optimization. However, we have found it sufficient to simply set *γ* to a small value (10^−3^ by default).

A simpler approach to optimization using the F1 distribution would be to simply run *K*(*K*+1)*/*2 different EM-algorithms — one for each possible combination of (*ℓ*_1_, *ℓ*_2_). However, we have found the approach of a single EM algorithm using a modified F1 distribution (39) much faster in practice and yields the same results. For example, on a 2.6 GHz quad-core PC running Linux with 32 GB of memory, running updog with the single EM approach with a discrete uniform mixing component took 2.6 seconds, 2.0 seconds, and 1.9 seconds on the three example SNPs used in Gerard et al. [2018]. On these same SNPs, running updog with the multiple EM approach took 15.7 seconds, 14.4 seconds, and 14.0 seconds. They yielded nearly the exact same posterior mean genotypes except in one individual in one SNP (Supplementary Figure S4).

To update the parameters from the class of unimodal distributions, we first assume that the “center” value of *a* is known (see Theorem 1). There exists a finite number of *a* values over which the likelihood varies which may be iterated over each M-step. Then let 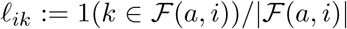. The prior optimization problem during the M-step then becomes

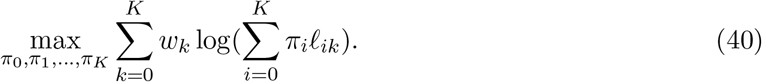

Equation (40) is a convex optimization problem and can be solved using standard interior-point methods. However, we found that in this particular problem (with a small value of *K*), we can obtain minor gains in performance by using a weighted EM algorithm to solve (40) (Supplementary Section S2.2).

### S2.2 Weighted EM derivation for the class of unimodal distributions

In this section, we describe a weighted EM algorithm to solve (40). This is an alternative optimization scheme to standard interior point methods. For the moment, suppose the *w*_*k*_’s in (40) are natural numbers. Then we can equivalently write (40) as

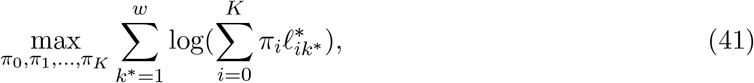

where

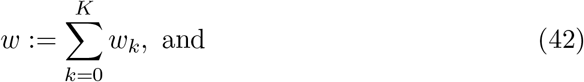

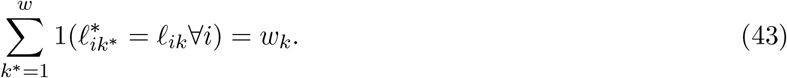

That is, for each *k* we make *w*_*k*_ copies of (*ℓ*_0*k*_, *ℓ*_1*k*_, … *ℓ*_*Kk*_). Equation (41) is equivalent to equation (S.2.6) of the Supplementary Material in Stephens [2016], and so the EM algorithm in Stephens [2016] may be used to update (41):

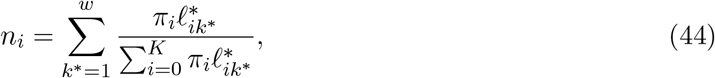

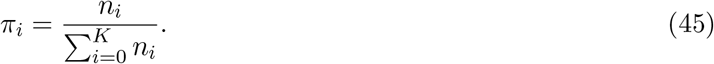

To solve (40) using the original parameter values, note that (44) is equivalent to

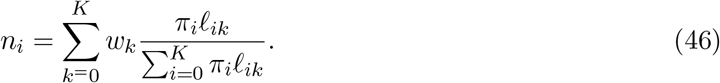

Since the update for the *π*_*i*_’s is invariant to scalar multiplication of the *w*_*k*_’s, we may update (40) for non-integer *w*_*k*_’s using (46) followed by (45).

The *ℓ*_*ik*_’s contain many exact zeros, which can cause convergence problem for the weighted EM algorithm. This can be remedied by placing a small Dirichlet prior on the *π*_*i*_’s (with the same concentration parameter placed on each *π*_*i*_). The resulting weighted EM algorithm is

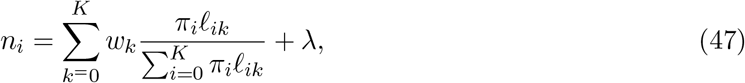

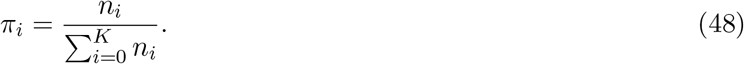

By default, we choose *λ* = 10^−6^.

We found that using the weighted EM algorithm performed faster in this scenario than using interior point methods. On a 2.6 GHz quad-core PC running Linux with 32 GB of memory, running updog with the weighted EM approach took 50.3 seconds, 34.4 seconds, 24.0 and seconds on the three example SNPs used in Gerard et al. [2018]. On these same SNPs, running updog with interior point methods (using the CVXR R package from Fu et al. [2017]) took 163.1 seconds, 87.8 seconds, and 66.2 seconds. The posterior mean genotypes were almost exactly the same using both optimization approaches (Supplementary Figure S5).

## S3 Genotype distributions used in the simulation study

In Section 3, genotypes were generated under the following seven conditions. Before each distribution, we list for which prior this genotype distribution was intended to be favorable.

1. Hardy-Weinberg Favorable: *Pr*(*k*) = Binomial(*k*|*K*, 0.75),
2. Beta-Binomial Favorable: *Pr*(*k*) = BB(*k*|*K*, 0.75, 0.1),
3. Proportional Normal Favorable: *Pr*(*k*) ∝ *N* (*k*|4.5, 0.5^2^),
4. Unimodal Favorable: 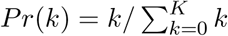,
5. F1 Favorable: *Pr*(*k*) was derived from one parent with 4 copies and one parent with 5 copies of reference allele,
6. General Favorable: ***π*** = (1, 4, 1, 4, 1, 4, 8)*/*23, and
7. Discrete Uniform Favorable: *Pr*(*k*) = 1*/*(*K* + 1).

These genotype distributions (except the discrete uniform distribution) are plotted in Supplementary Figure S6.

## S4

**Figure S1:**
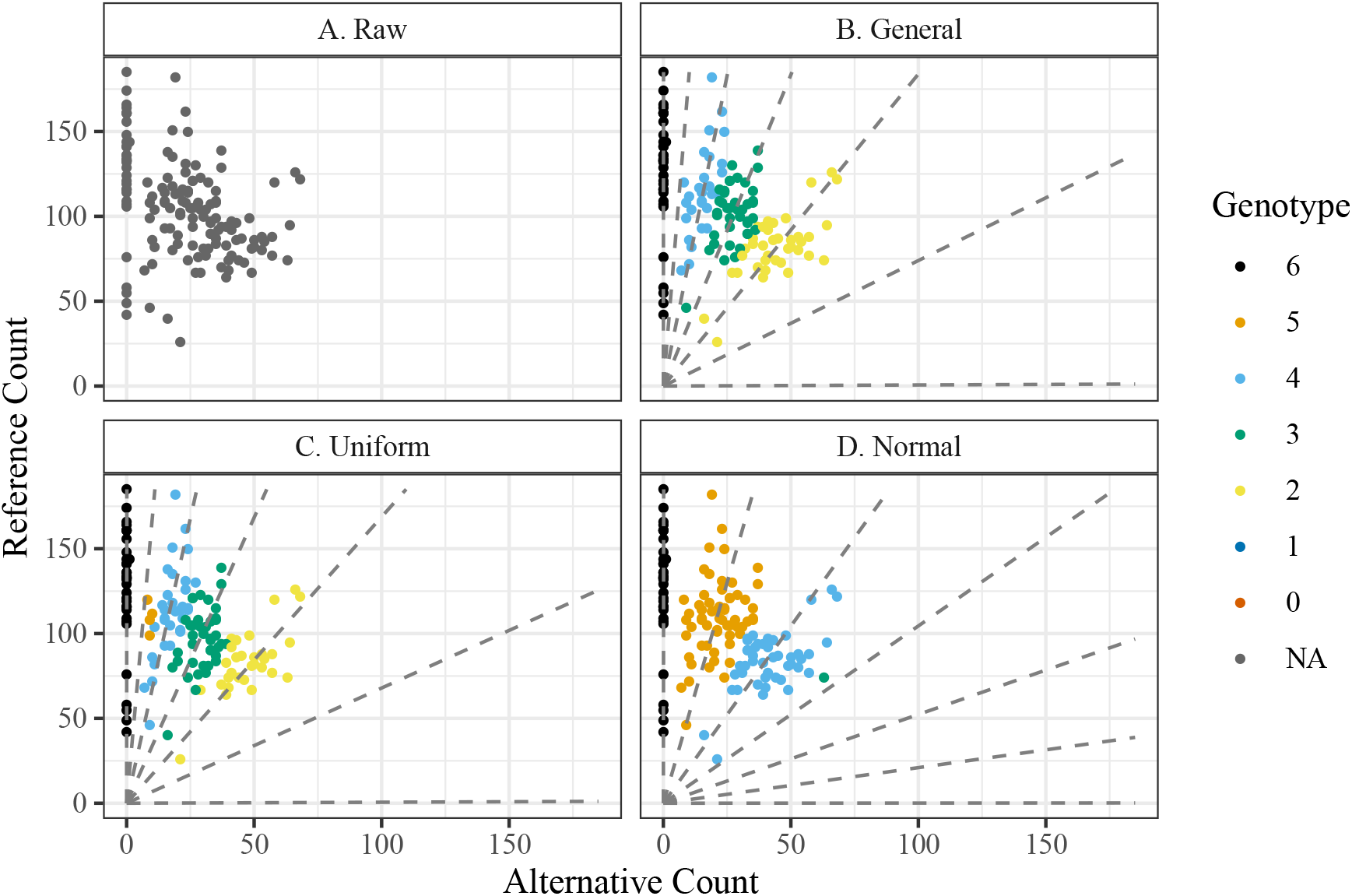
The number of reference reads (*y*-axis) versus the number of alternative reads (*x*-axis) for a SNP from 143 sweet potatoes from Shirasawa et al. [2017]. The raw unannotated data are presented in panel (A). The other panels are color-coded by estimated genotype according to updog fits using either (B) the general class of priors, (C) the discrete uniform prior distribution, or (D) the class of proportional normal priors. The dashed lines in panels (B) through (D) are the estimated expected counts, and are functions of the estimated allele bias and estimated sequencing error rate [see Gerard et al., 2018]. These individuals are known to be siblings resulting from an S1 cross. Given this knowledge, only the proportional normal class of priors provides intuitive genotyping.

**Table S1:** Parameter values for the proportional normal distributions displayed in Supplementary Figure S3.

| Distribution | Distribution Parameters | Normal Mean | Normal SD |
| --- | --- | --- | --- |
| Uniform | · | 3 | ∞ |
| F1 | $\ell_1 = 1$ and $\ell_2 = 2$ | 1.5 | 0.8 |
| Hardy-Weinberg (Binomial) | $\alpha = 0.25$ | 1.4 | 1.2 |
| Beta-binomial | $\alpha = 0.25$ and $\gamma = 0.01$ | 0.8 | 1.8 |

**Figure S2:**
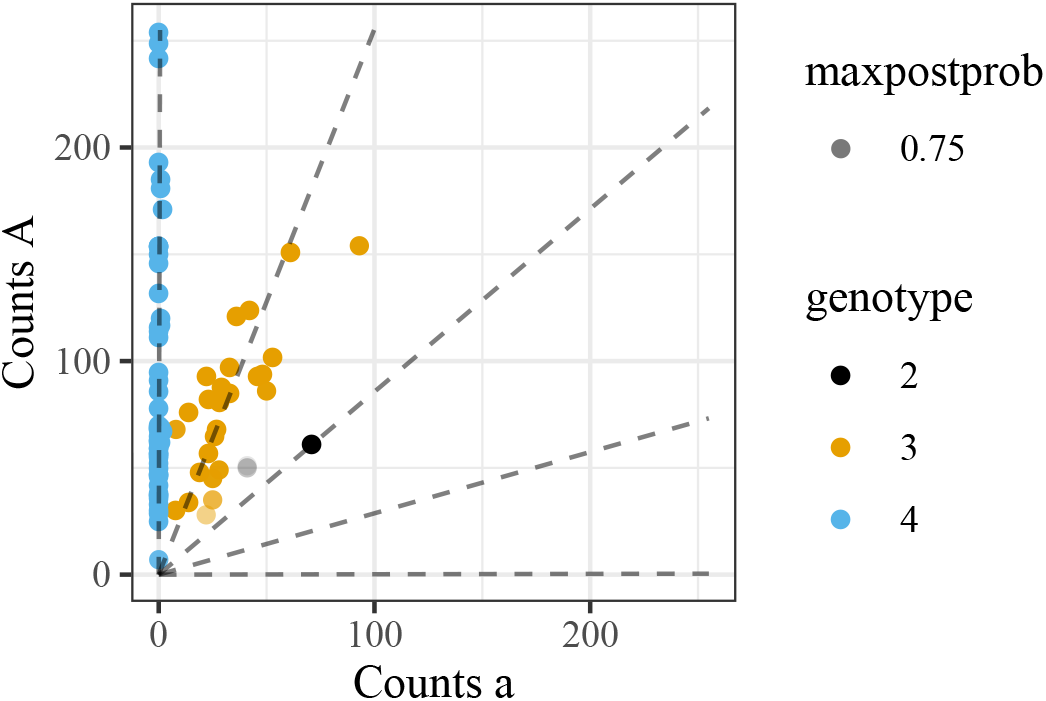
An annotated genotype plot, same as panel (B) of Figure 1 except a stronger penalty was applied to the bias parameter during the updog fit. This strengthened penalty resulted in more intuitive genotyping.

**Table S2:**
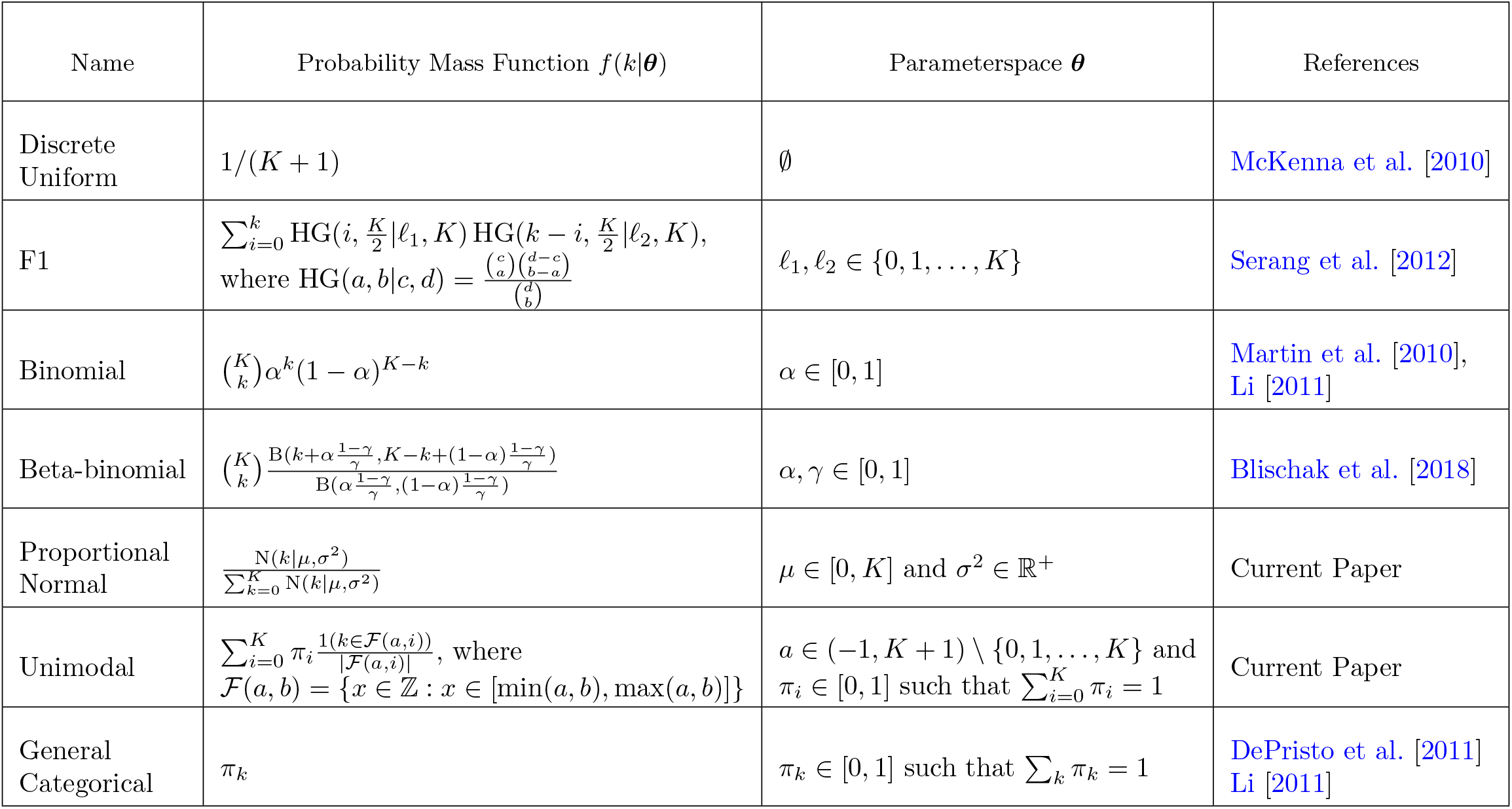
Prior distributions considered in this paper. We list their names, probability mass functions, the parameterspaces of the prior-specific parameters ***θ***, and references where these priors were used for genotyping.

**Figure S3:**
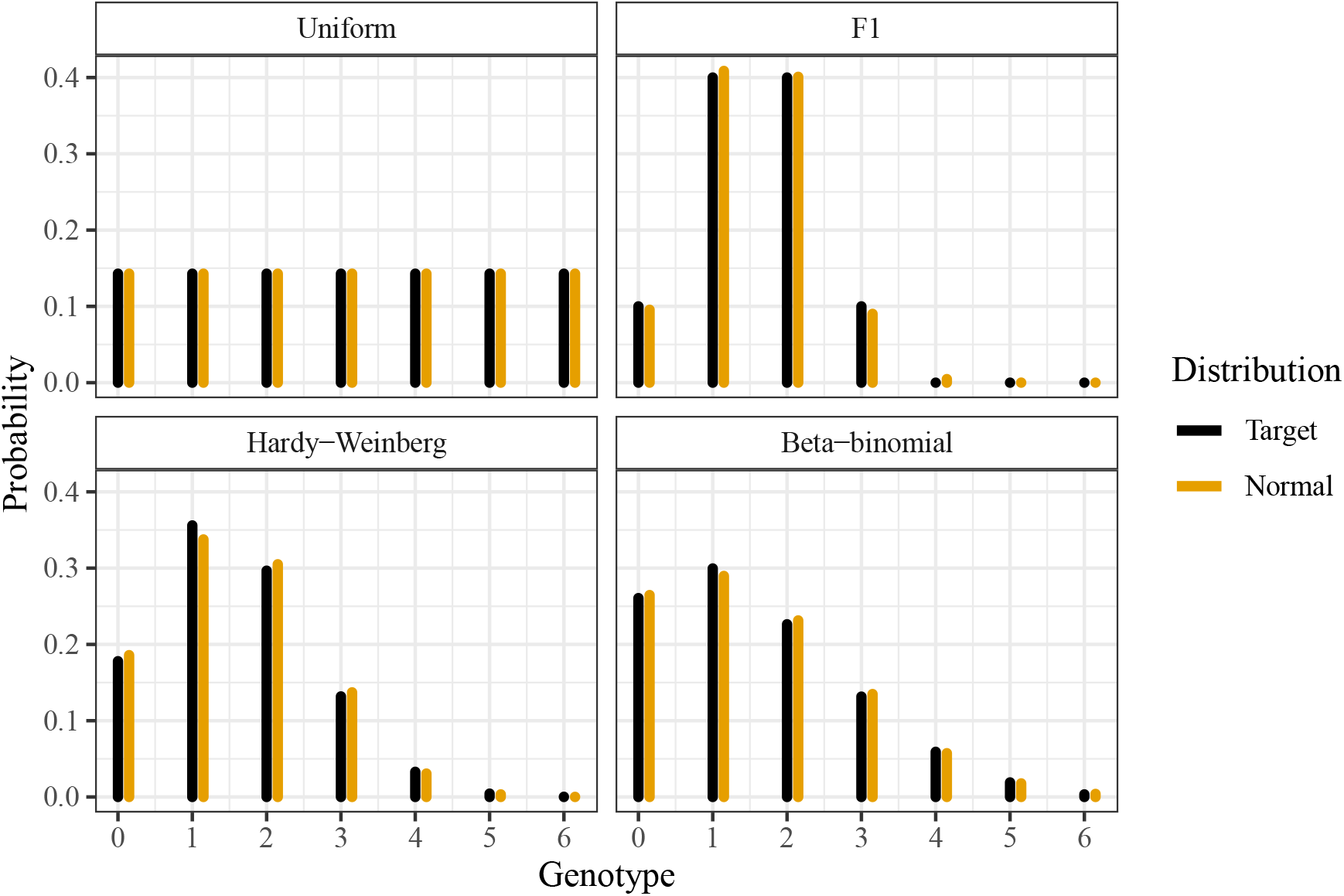
Four distributions of genotypes (black) and the closest distribution in the class of proportional normal distributions (orange). Closeness was measured by Kullback–Leibler divergence. Elements in the class of proportional normal distributions approximate very well the discrete uniform, the F1 (*ℓ*_1_ = 1 and *ℓ*_2_ = 2), the binomial (*α* = 0.25), and the beta-binomial (*α* = 0.25 and *γ* = 0.01) distributions. The parameter values of the plotted proportional normal distributions are given in Supplementary Table S1.

**Figure S4:**
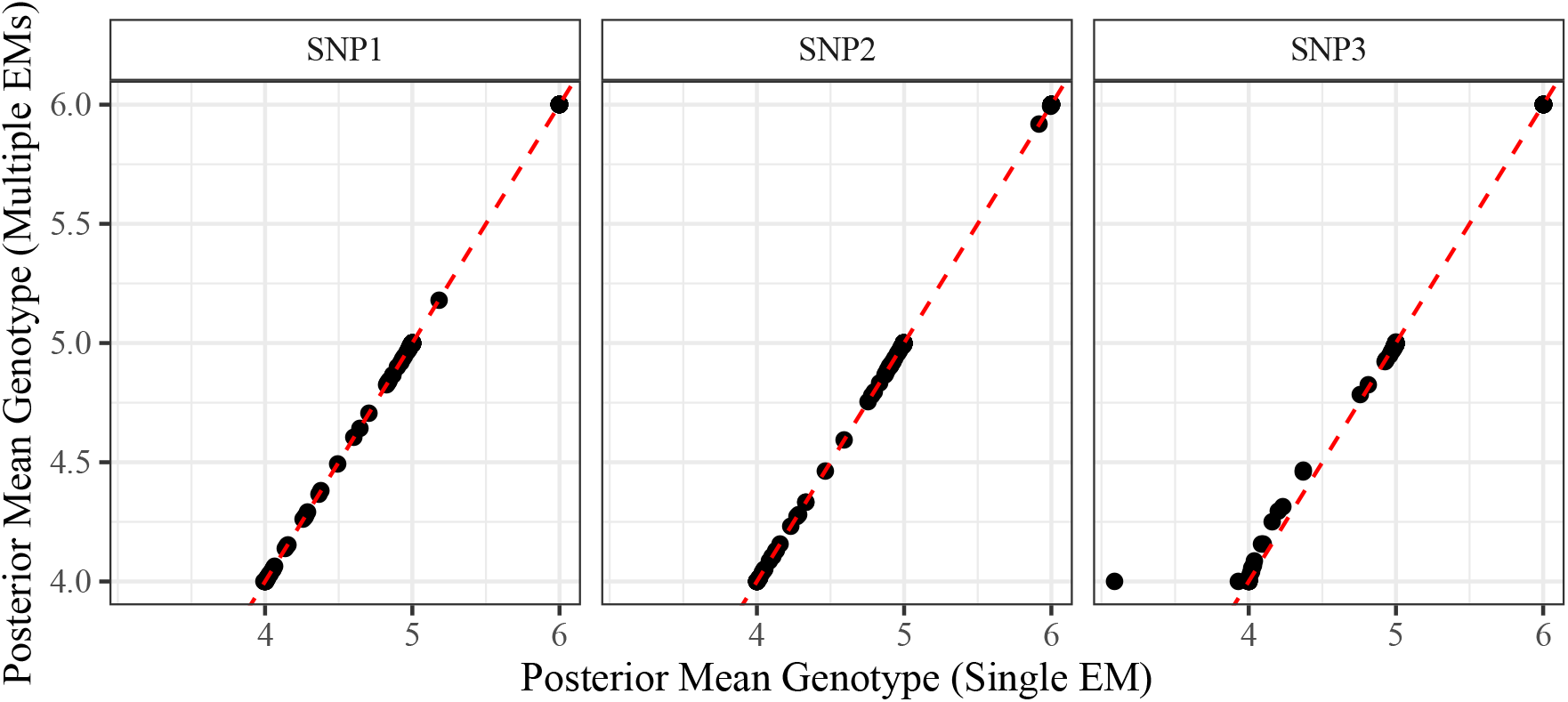
Posterior mean genotypes for the three example SNPs used in Gerard et al. [2018]. These were calculated using either the single EM approach with a small discrete uniform mixing component (*x*-axis) or the multiple EM approach that iterates over all possible parental genotypes (*y*-axis). The dashed red line is the *y* = *x* line. The posterior mean genotypes from the two approaches are nearly identical.

**Figure S5:**
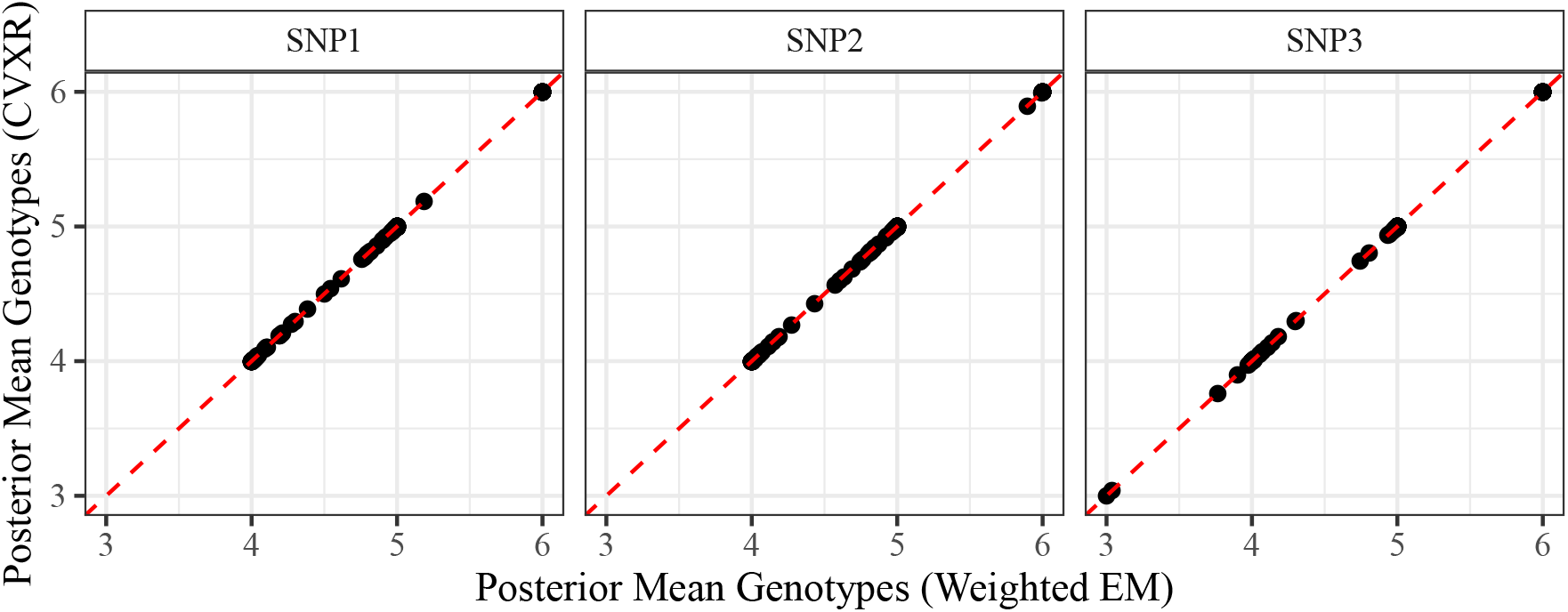
Posterior mean genotypes for the three example SNPs used in Gerard et al. [2018]. These were calculated using either the weighted EM approach for the class of unimodal distributions (*x*-axis) or using interior point methods (*y*-axis). The dashed red line is the *y* = *x* line. The posterior mean genotypes from the two approaches are nearly identical.

**Figure S6:**
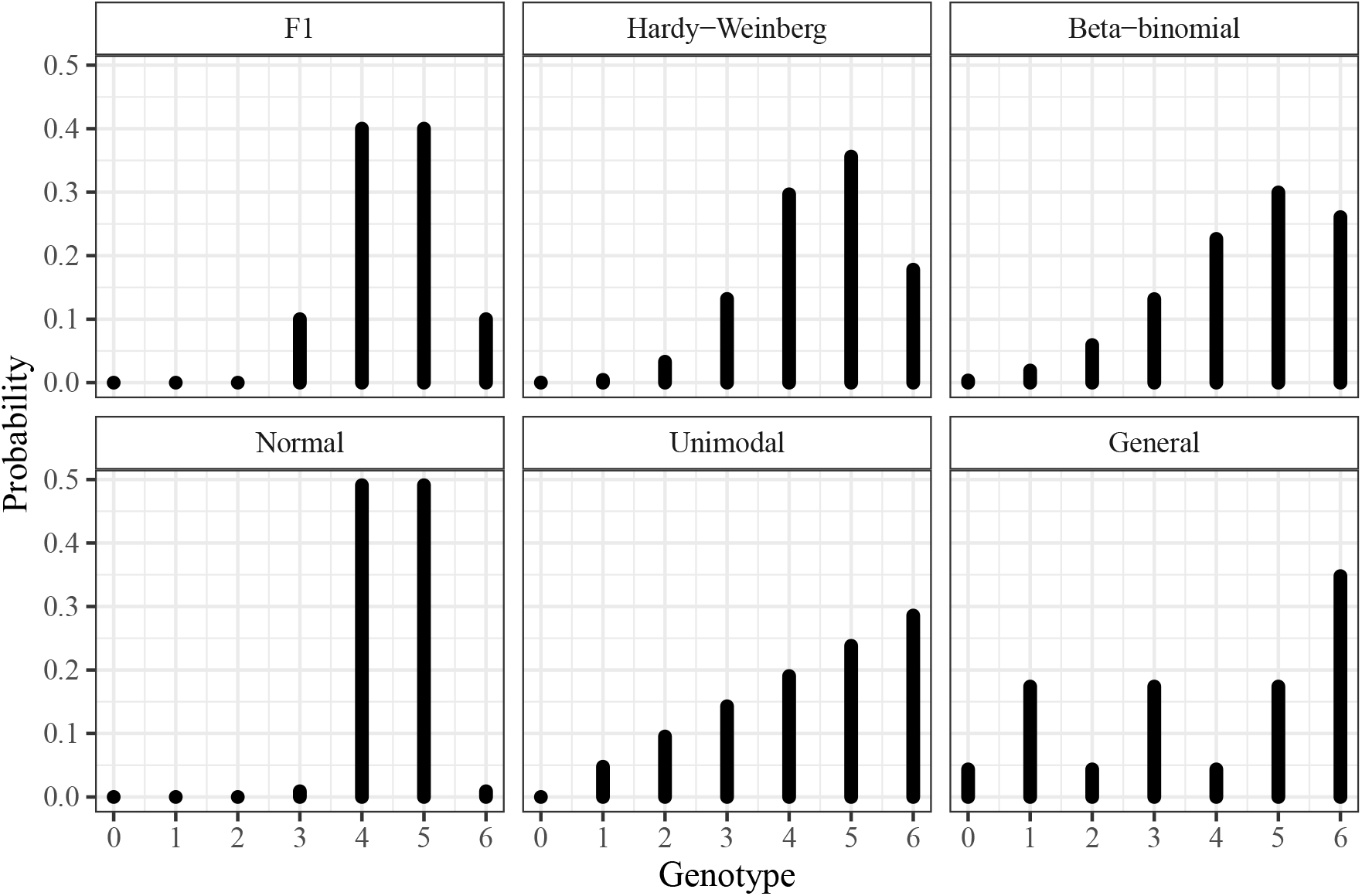
True genotype distributions used in the simulation study of Section 3, explicitly defined in Supplementary Section S3. Each genotype distribution was chosen to be favorable to one of the assumed classes of priors: the class of F1 distributions, the class of binomial (Hardy-Weinberg) distributions, the class of beta-binomial distributions, the class of proportional normal distributions, the class of unimodal distributions, and the class of general distributions. Not plotted is the discrete uniform distribution, also used in the simulation study.

**Figure S7:**
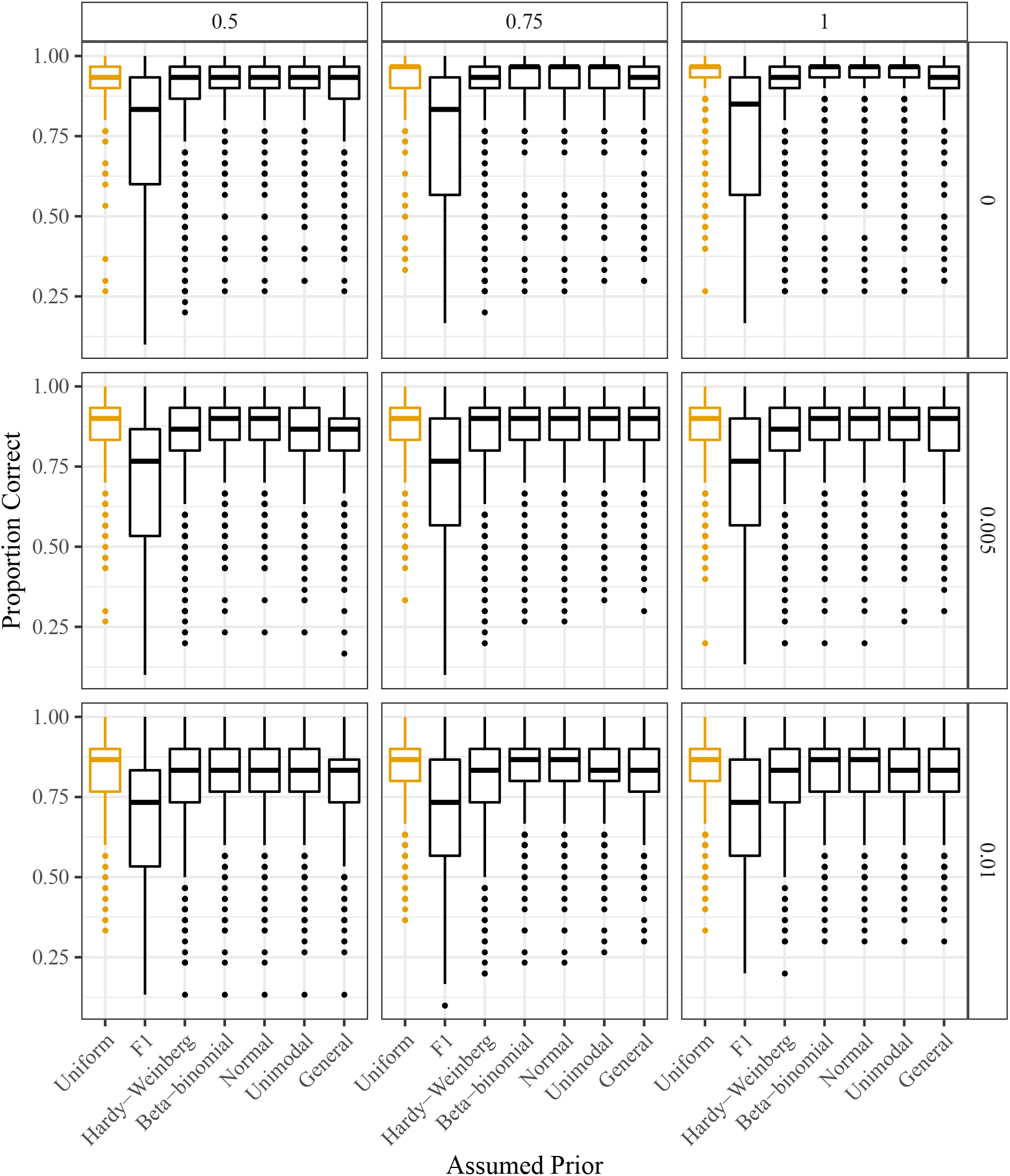
Proportion of individuals genotyped correctly (*y*-axis) stratified by the assumed class of prior distribution (*x*-axis). Column facets index varying levels of allele bias (with 1 indicating no bias) and row facets index varying levels of overdispersion (with 0 indicating no overdispersion). The genotypes were generated from a distribution designed to be favorable to the discrete uniform prior distribution (orange).

**Figure S8:**
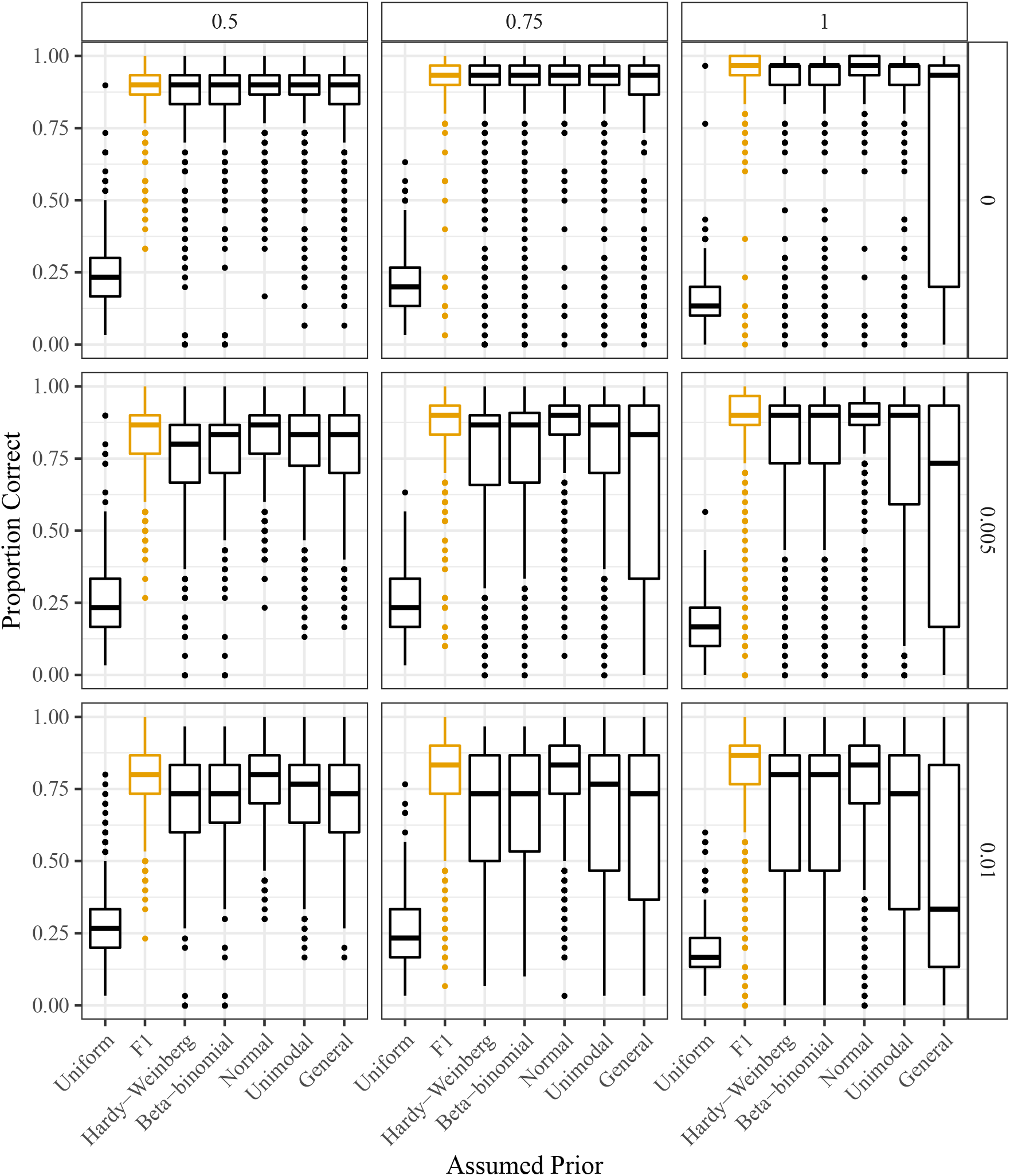
Proportion of individuals genotyped correctly (*y*-axis) stratified by the assumed class of prior distribution (*x*-axis). Column facets index varying levels of allele bias (with 1 indicating no bias) and row facets index varying levels of overdispersion (with 0 indicating no overdispersion). The genotypes were generated from a distribution designed to be favorable to the F1 class of prior distributions (orange).

**Figure S9:**
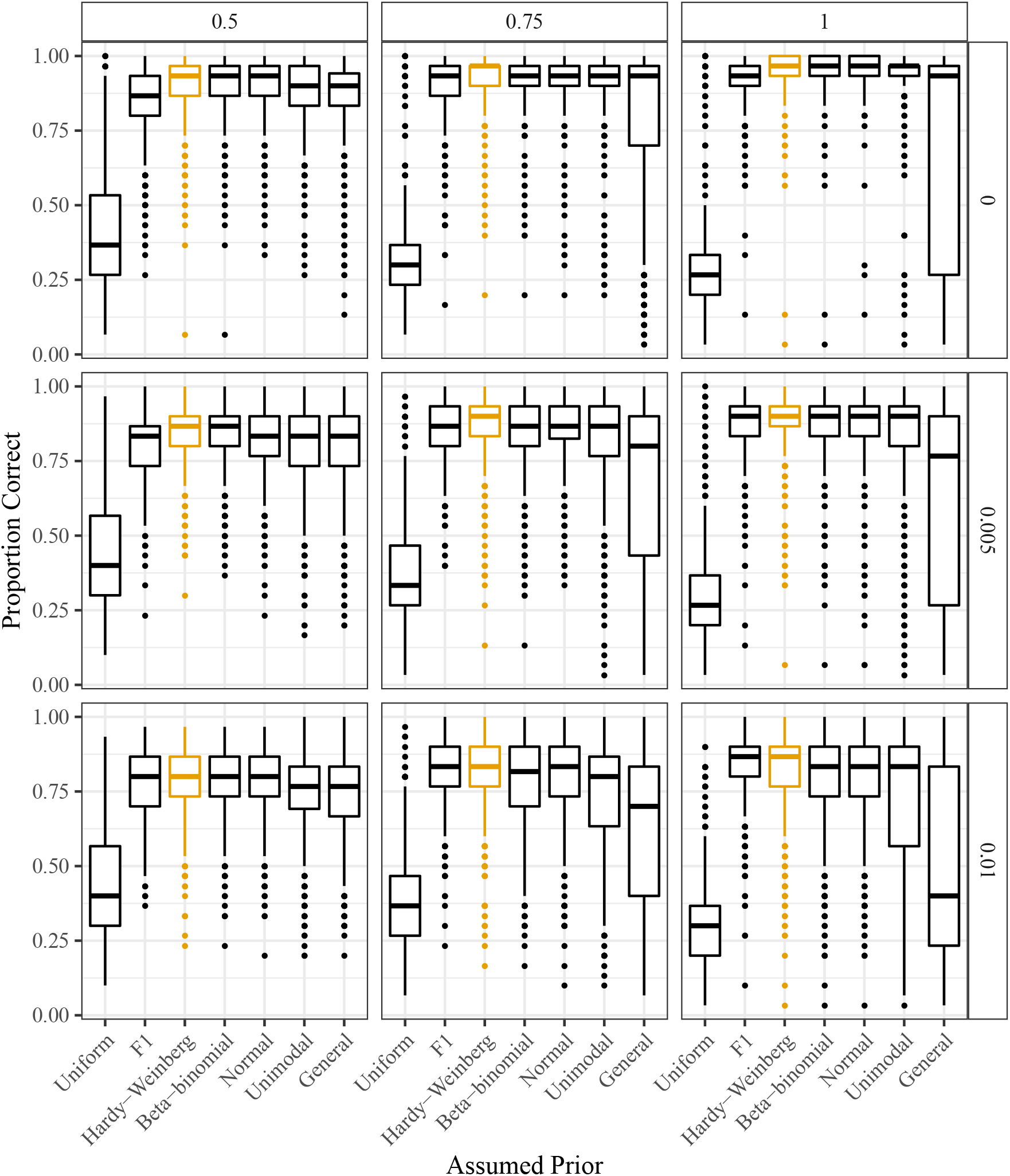
Proportion of individuals genotyped correctly (*y*-axis) stratified by the assumed class of prior distribution (*x*-axis). Column facets index varying levels of allele bias (with 1 indicating no bias) and row facets index varying levels of overdispersion (with 0 indicating no overdispersion). The genotypes were generated from a distribution designed to be favorable to the binomial (Hardy-Weinberg) class of prior distributions (orange).

**Figure S10:**
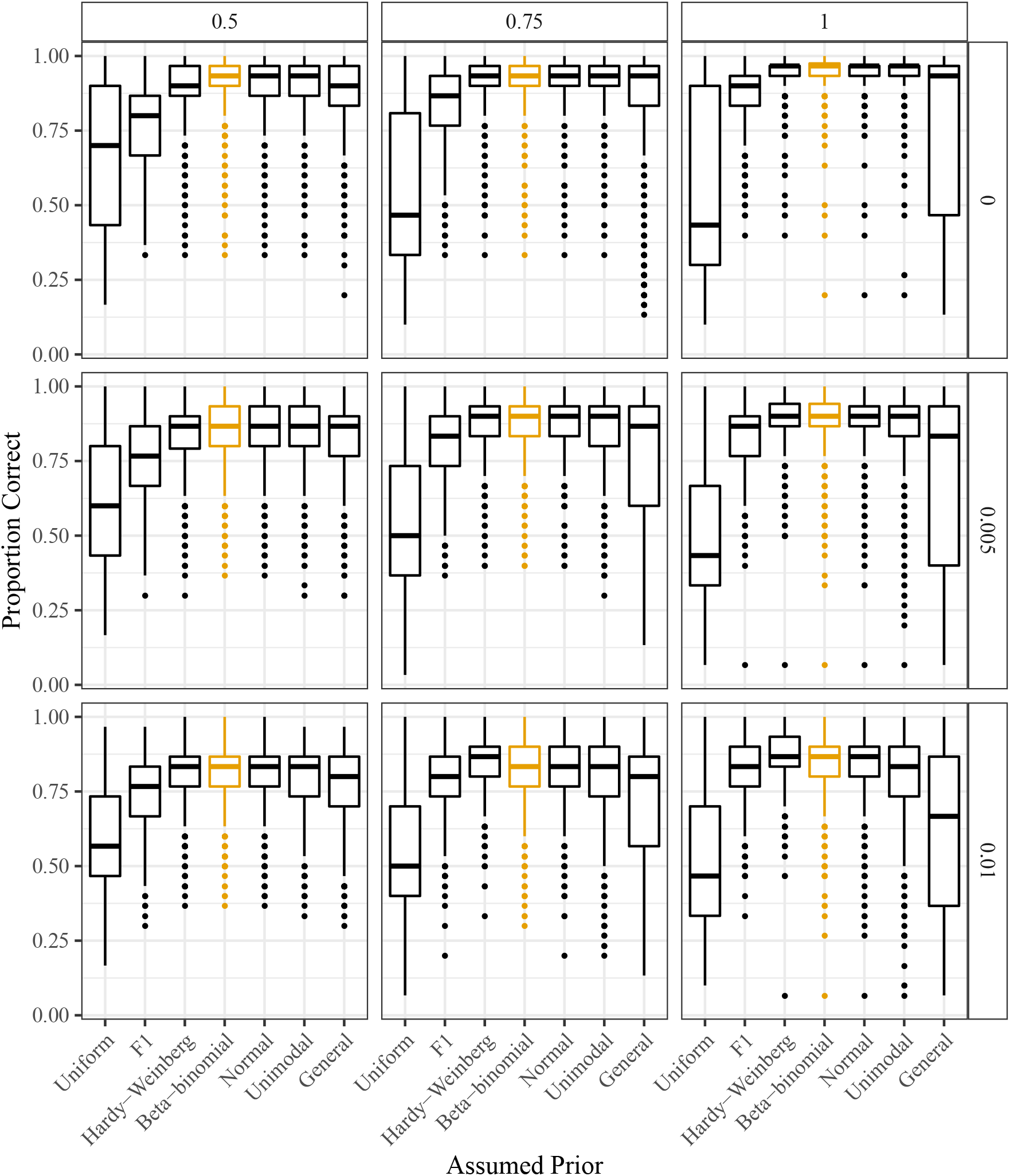
Proportion of individuals genotyped correctly (*y*-axis) stratified by the assumed class of prior distribution (*x*-axis). Column facets index varying levels of allele bias (with 1 indicating no bias) and row facets index varying levels of overdispersion (with 0 indicating no overdispersion). The genotypes were generated from a distribution designed to be favorable to the beta-binomial class of prior distributions (orange).

**Figure S11:**
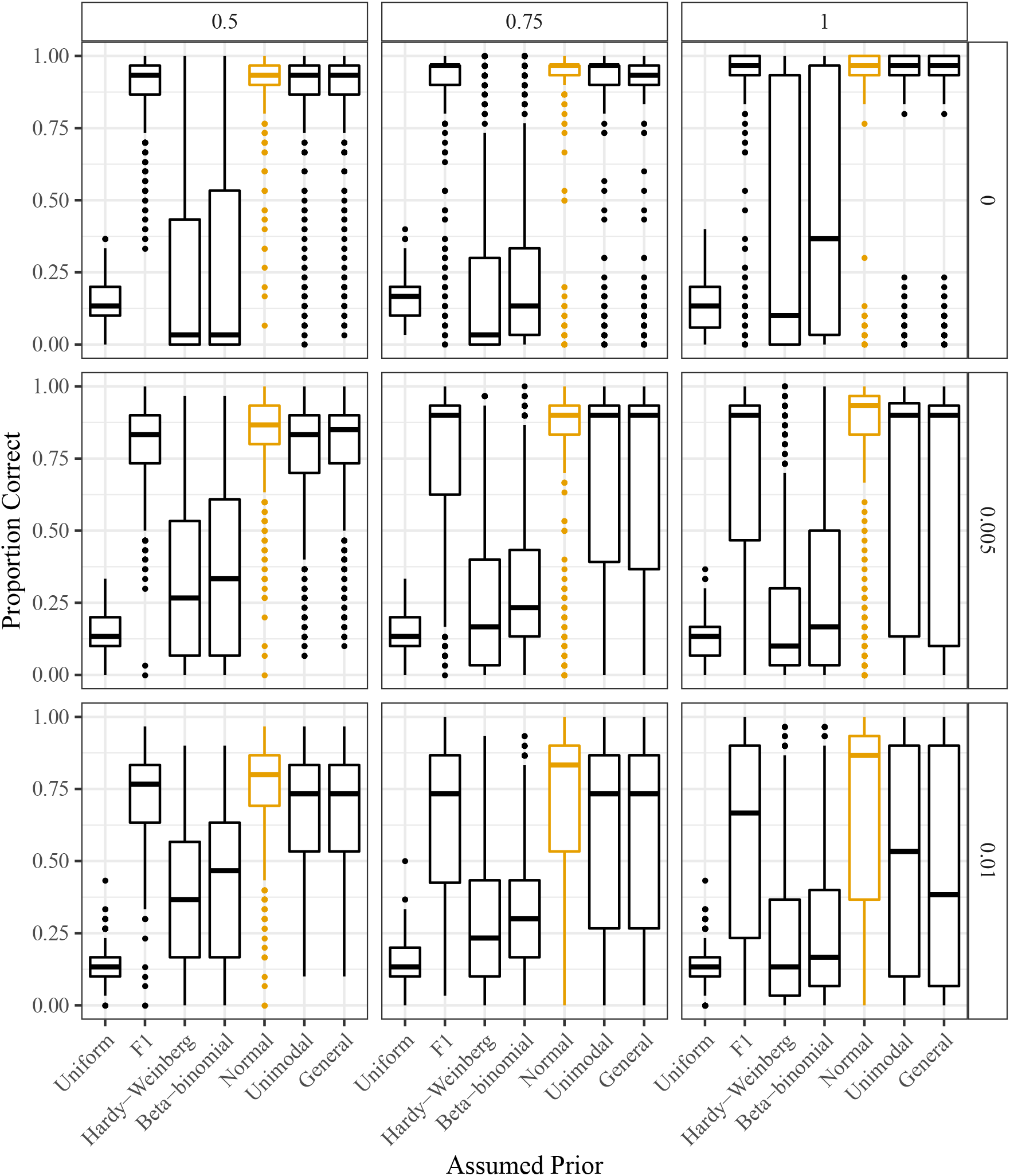
Proportion of individuals genotyped correctly (*y*-axis) stratified by the assumed class of prior distribution (*x*-axis). Column facets index varying levels of allele bias (with 1 indicating no bias) and row facets index varying levels of overdispersion (with 0 indicating no overdispersion). The genotypes were generated from a distribution designed to be favorable to the proportional normal class of prior distributions (orange).

**Figure S12:**
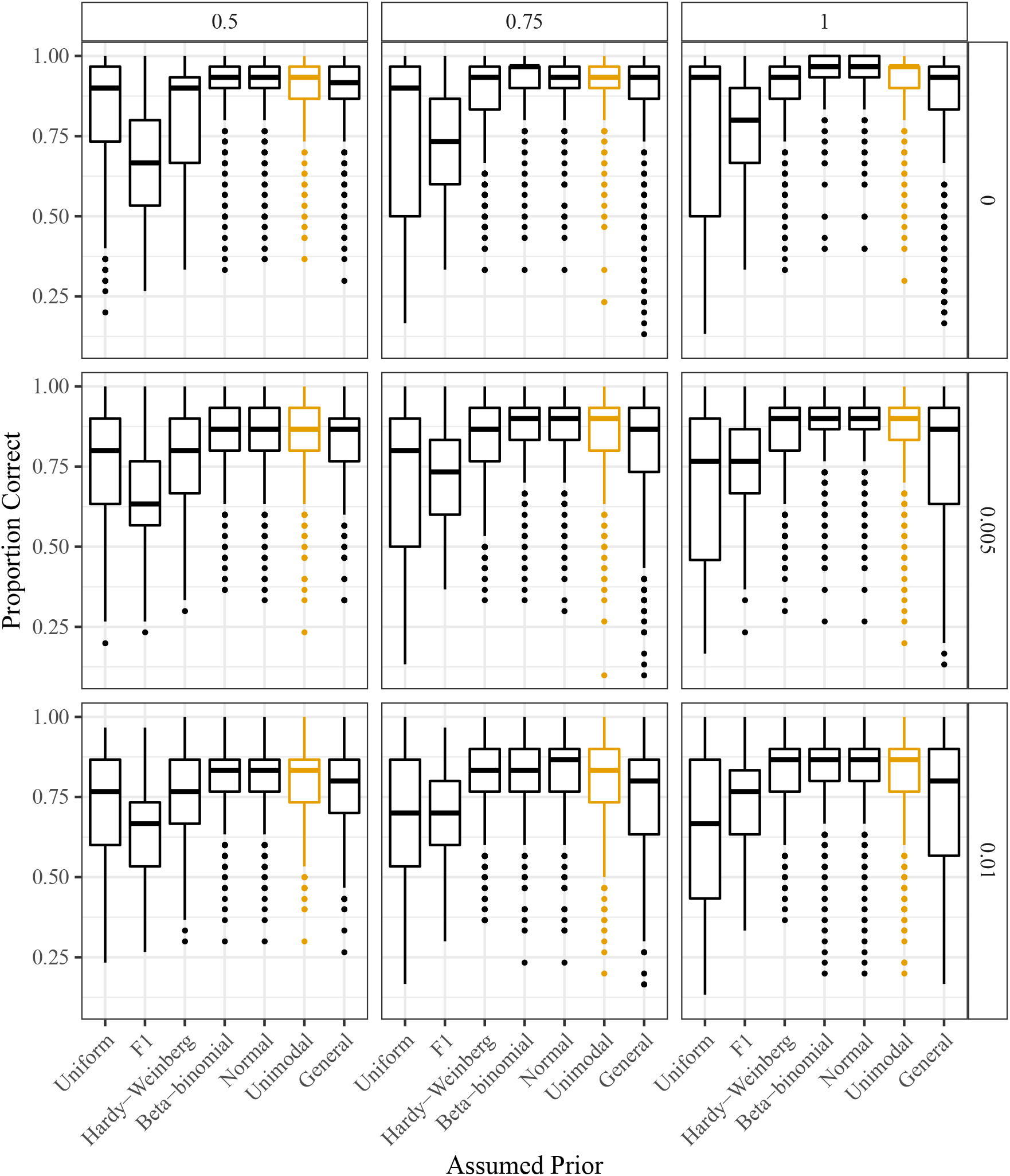
Proportion of individuals genotyped correctly (*y*-axis) stratified by the assumed class of prior distribution (*x*-axis). Column facets index varying levels of allele bias (with 1 indicating no bias) and row facets index varying levels of overdispersion (with 0 indicating no overdispersion). The genotypes were generated from a distribution designed to be favorable to the unimodal class of prior distributions (orange).

**Figure S13:**
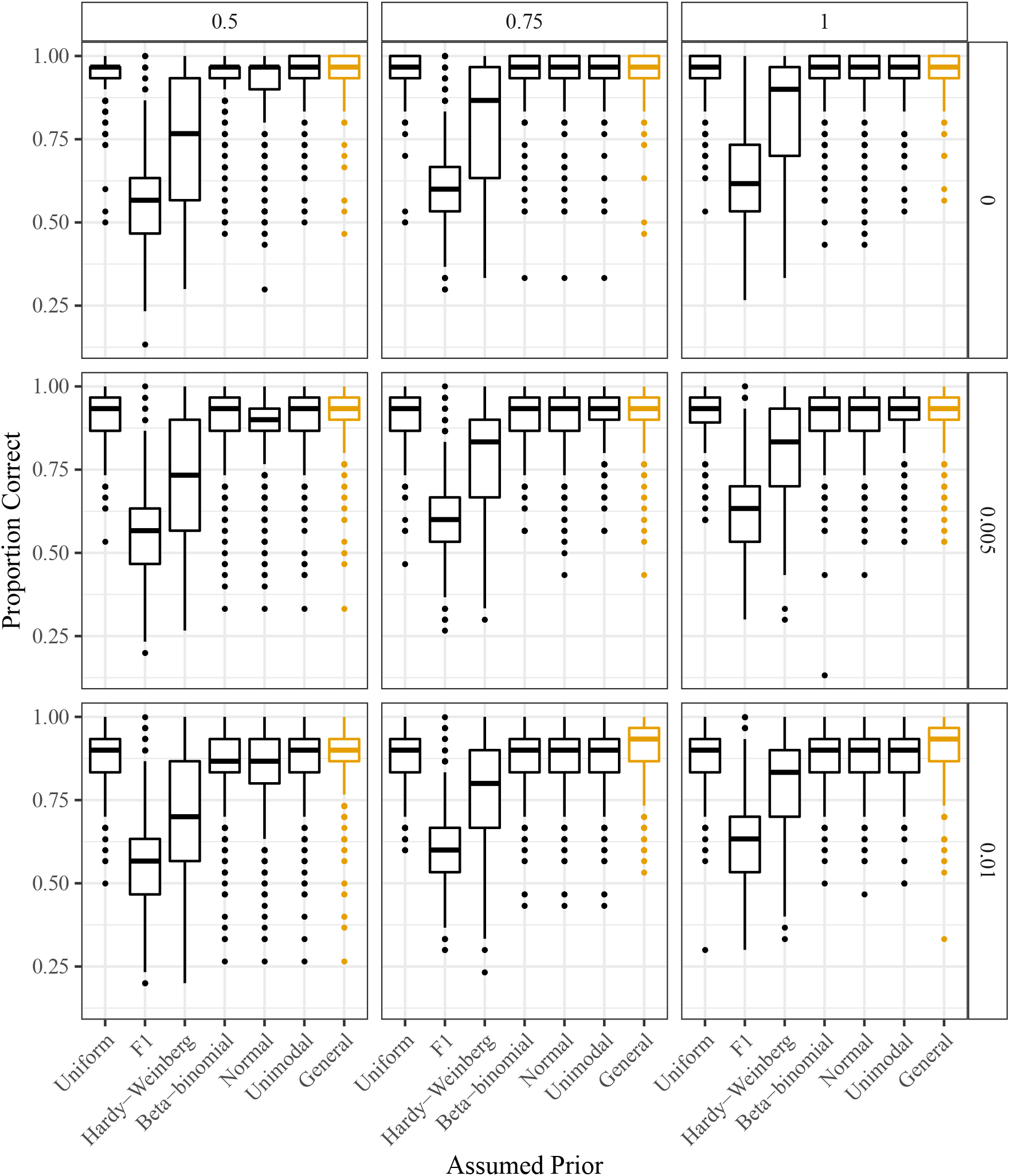
Proportion of individuals genotyped correctly (*y*-axis) stratified by the assumed class of prior distributions (*x*-axis). Column facets index varying levels of allele bias (with 1 indicating no bias) and row facets index varying levels of overdispersion (with 0 indicating no overdispersion). The genotypes were generated from a distribution designed to be favorable to the general class of prior distributions (orange).

**Figure S14:**
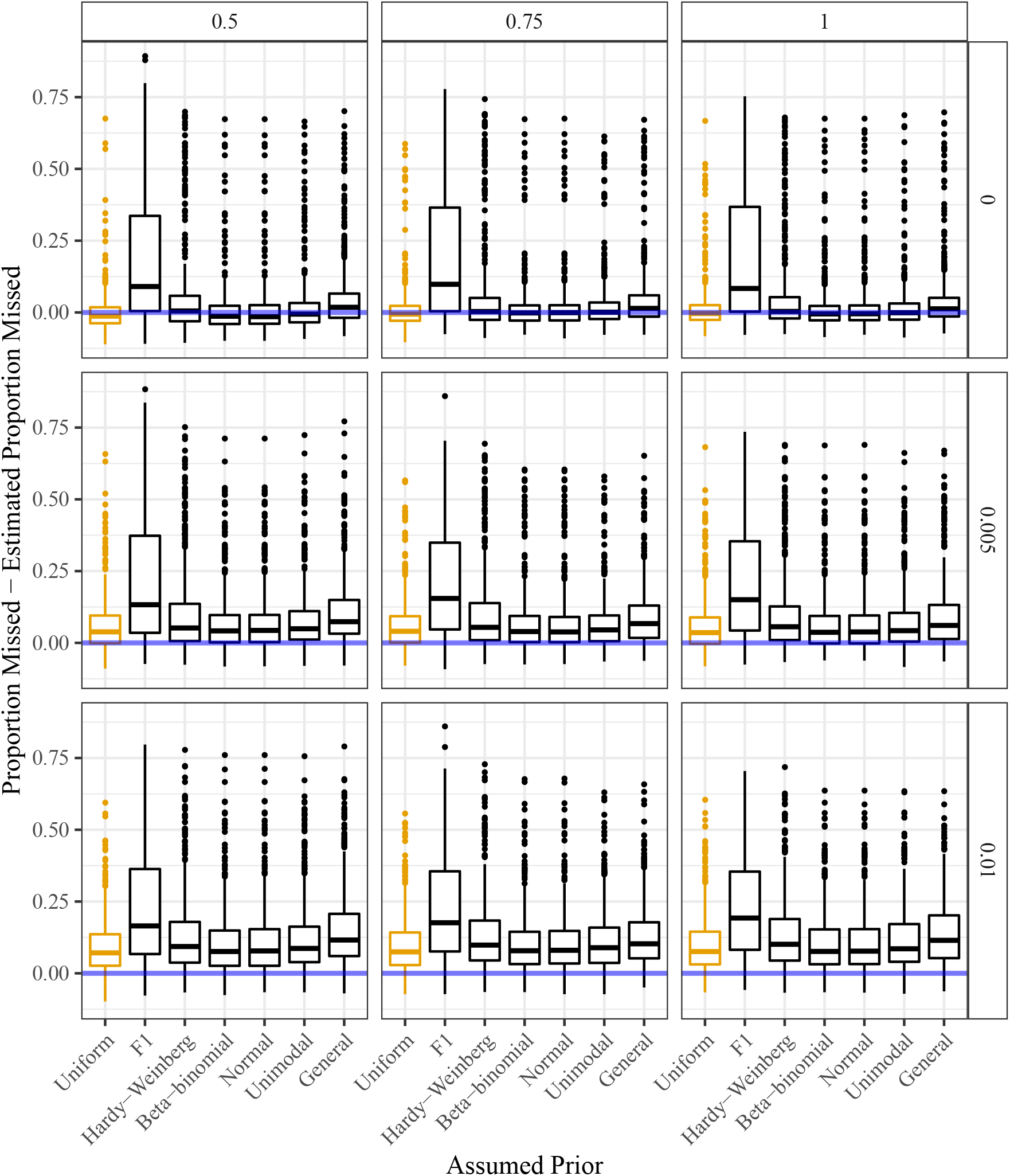
Estimated proportion of individuals incorrectly genotyped subtracted from the actual proportion of individuals incorrectly genotyped (*y*-axis) stratified by the assumed class of prior distributions (*x*-axis). The horizontal line (at *y* = 0) indicates unbiased estimation of the misclassification error rate. Column facets index varying levels of allele bias (with 1 indicating no bias) and row facets index varying levels of overdispersion (with 0 indicating no overdispersion). The genotypes were generated from a distribution designed to be favorable to the discrete uniform prior distribution (orange).

**Figure S15:**
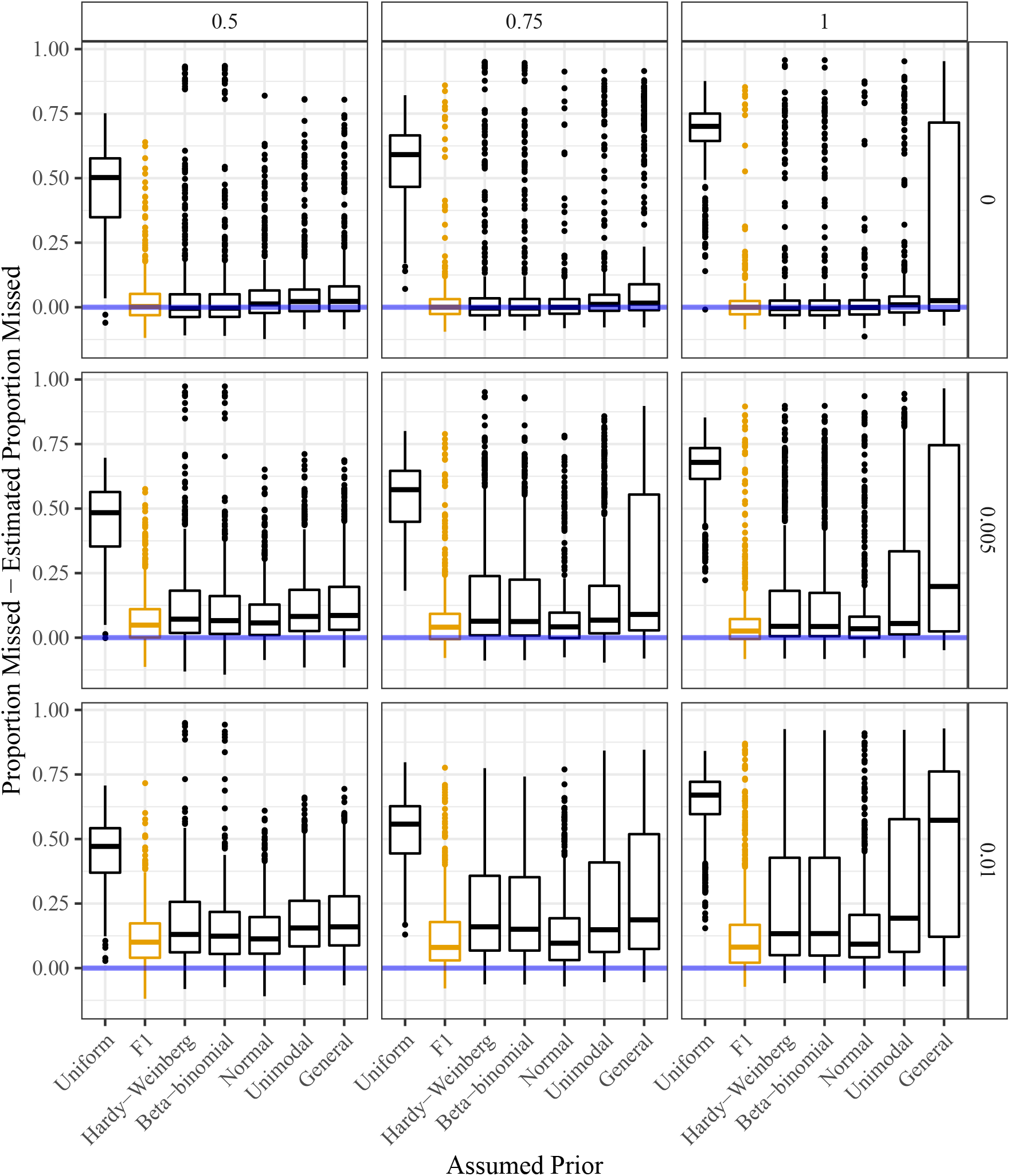
Estimated proportion of individuals incorrectly genotyped subtracted from the actual proportion of individuals incorrectly genotyped (*y*-axis) stratified by the assumed class of prior distributions (*x*-axis). The horizontal line (at *y* = 0) indicates unbiased estimation of the misclassification error rate. Column facets index varying levels of allele bias (with 1 indicating no bias) and row facets index varying levels of overdispersion (with 0 indicating no overdispersion). The genotypes were generated from a distribution designed to be favorable to the class of F1 prior distributions (orange).

**Figure S16:**
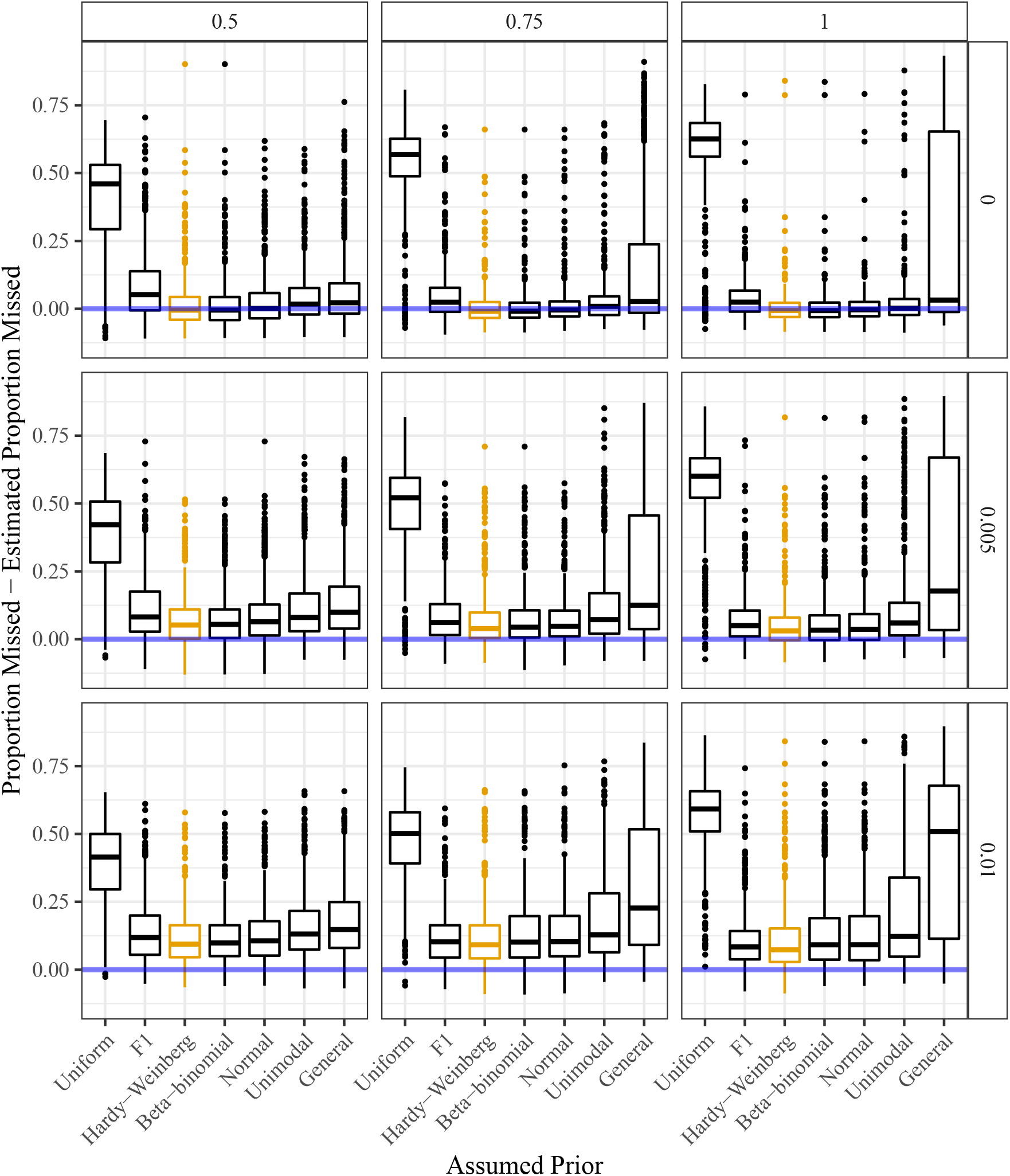
Estimated proportion of individuals incorrectly genotyped subtracted from the actual proportion of individuals incorrectly genotyped (*y*-axis) stratified by the assumed class of prior distributions (*x*-axis). The horizontal line (at *y* = 0) indicates unbiased estimation of the misclassification error rate. Column facets index varying levels of allele bias (with 1 indicating no bias) and row facets index varying levels of overdispersion (with 0 indicating no overdispersion). The genotypes were generated from a distribution designed to be favorable to the class of binomial (Hardy-Weinberg) prior distributions (orange).

**Figure S17:**
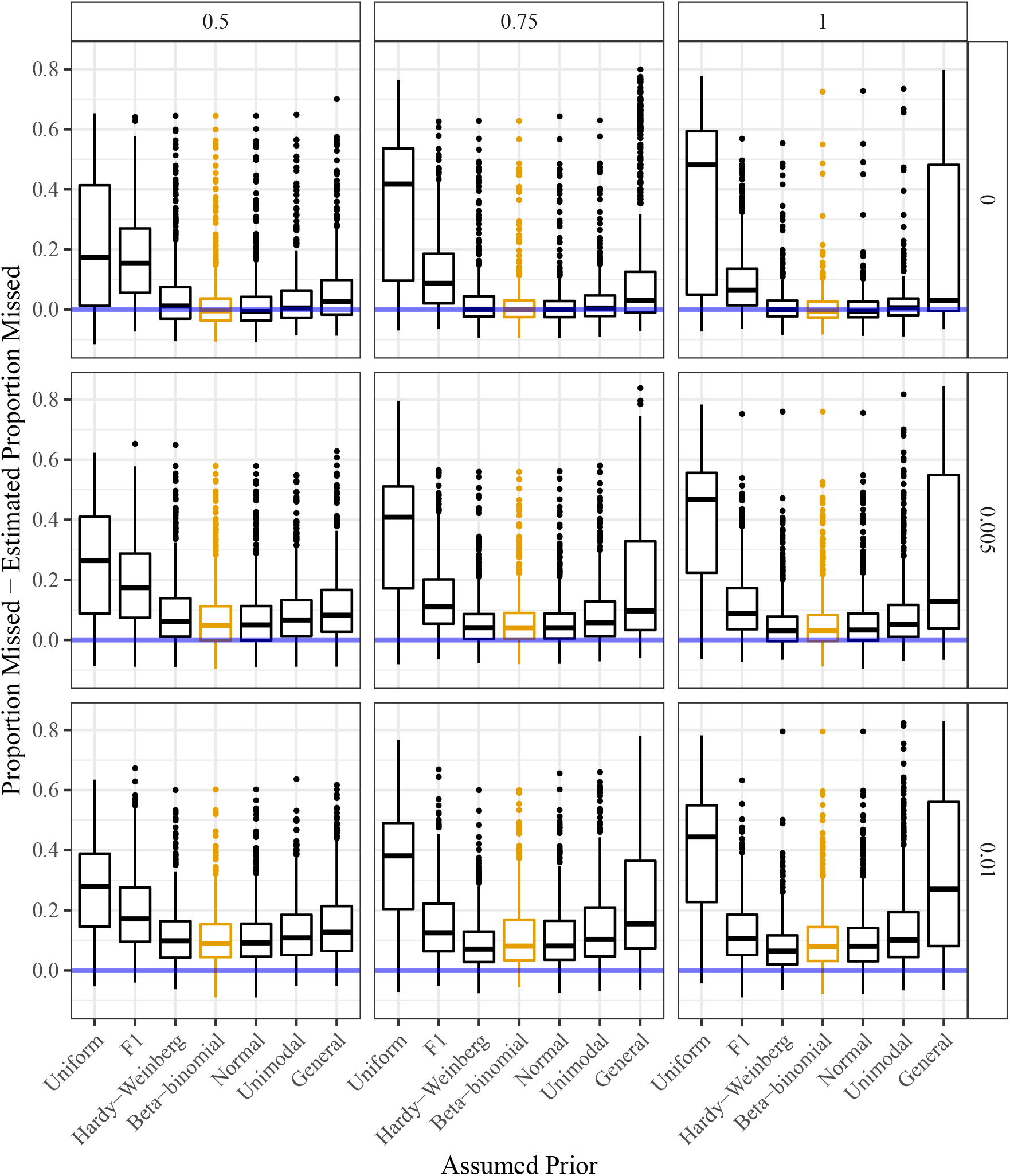
Estimated proportion of individuals incorrectly genotyped subtracted from the actual proportion of individuals incorrectly genotyped (*y*-axis) stratified by the assumed class of prior distributions (*x*-axis). The horizontal line (at *y* = 0) indicates unbiased estimation of the misclassification error rate. Column facets index varying levels of allele bias (with 1 indicating no bias) and row facets index varying levels of overdispersion (with 0 indicating no overdispersion). The genotypes were generated from a distribution designed to be favorable to the class of beta-binomial prior distributions (orange).

**Figure S18:**
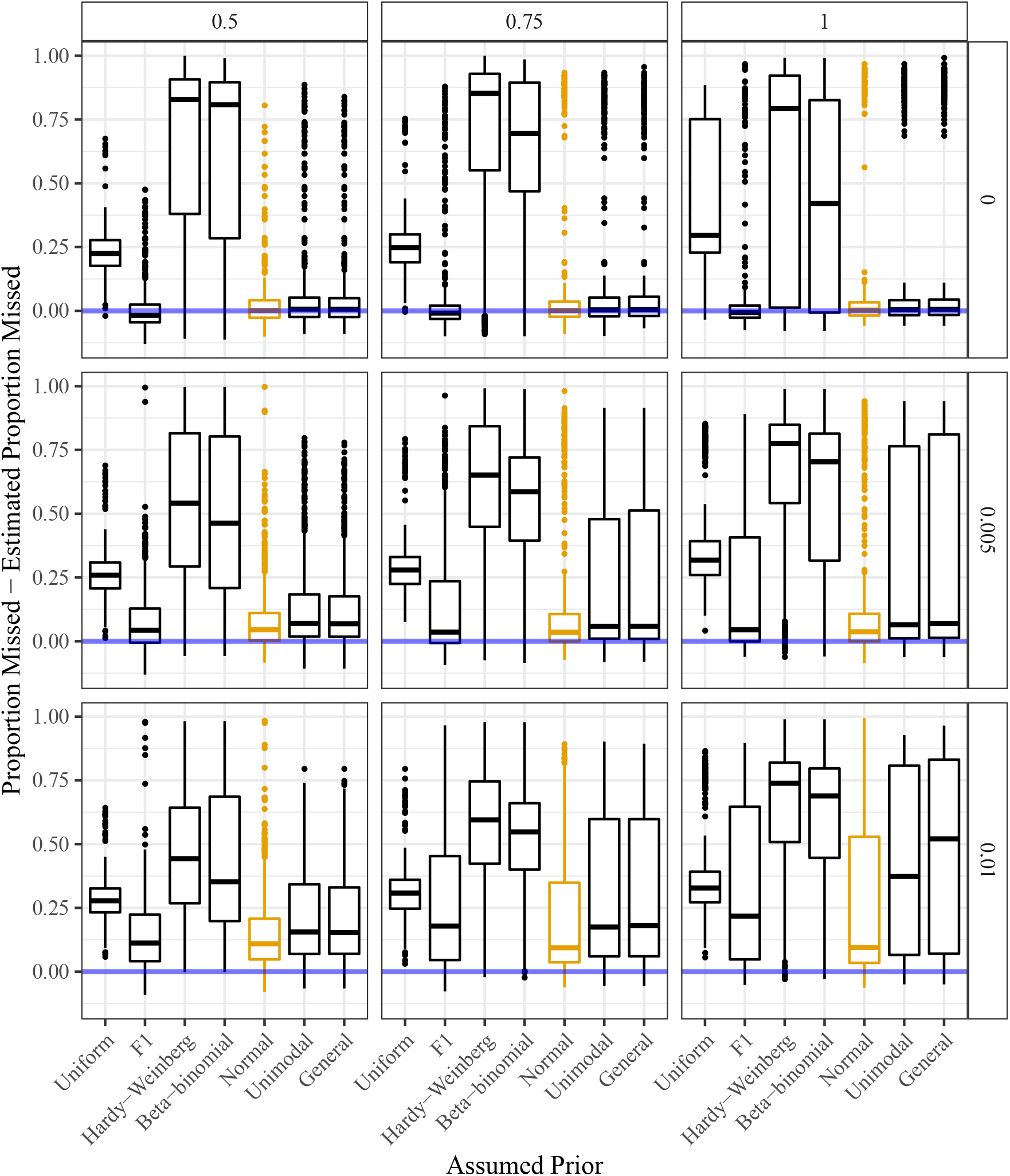
Estimated proportion of individuals incorrectly genotyped subtracted from the actual proportion of individuals incorrectly genotyped (*y*-axis) stratified by the assumed class of prior distributions (*x*-axis). The horizontal line (at *y* = 0) indicates unbiased estimation of the misclassification error rate. Column facets index varying levels of allele bias (with 1 indicating no bias) and row facets index varying levels of overdispersion (with 0 indicating no overdispersion). The genotypes were generated from a distribution designed to be favorable to the class of proportional normal prior distributions (orange).

**Figure S19:**
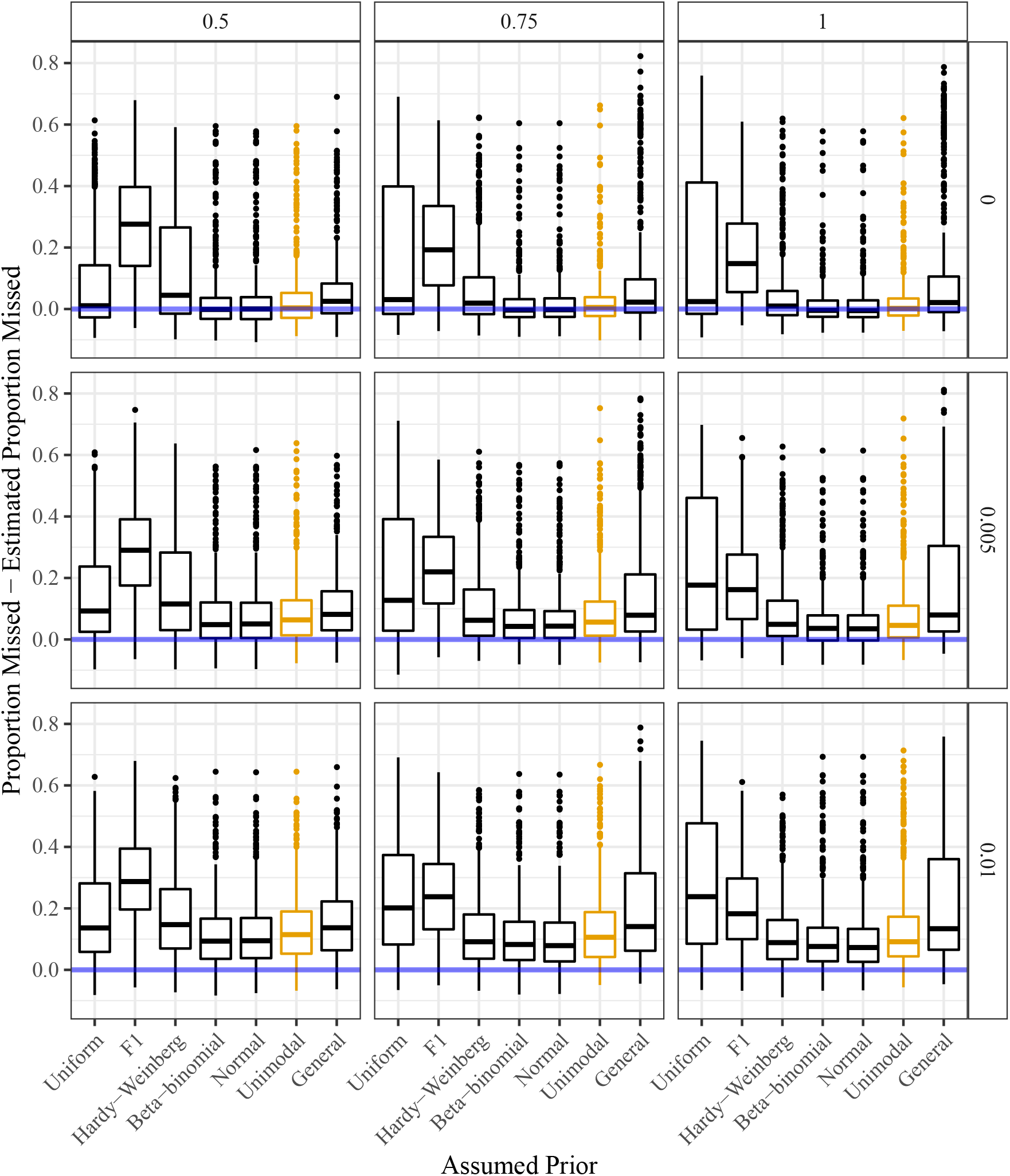
Estimated proportion of individuals incorrectly genotyped subtracted from the actual proportion of individuals incorrectly genotyped (*y*-axis) stratified by the assumed class of prior distributions (*x*-axis). The horizontal line (at *y* = 0) indicates unbiased estimation of the misclassification error rate. Column facets index varying levels of allele bias (with 1 indicating no bias) and row facets index varying levels of overdispersion (with 0 indicating no overdispersion). The genotypes were generated from a distribution designed to be favorable to the class of unimodal prior distributions (orange).

**Figure S20:**
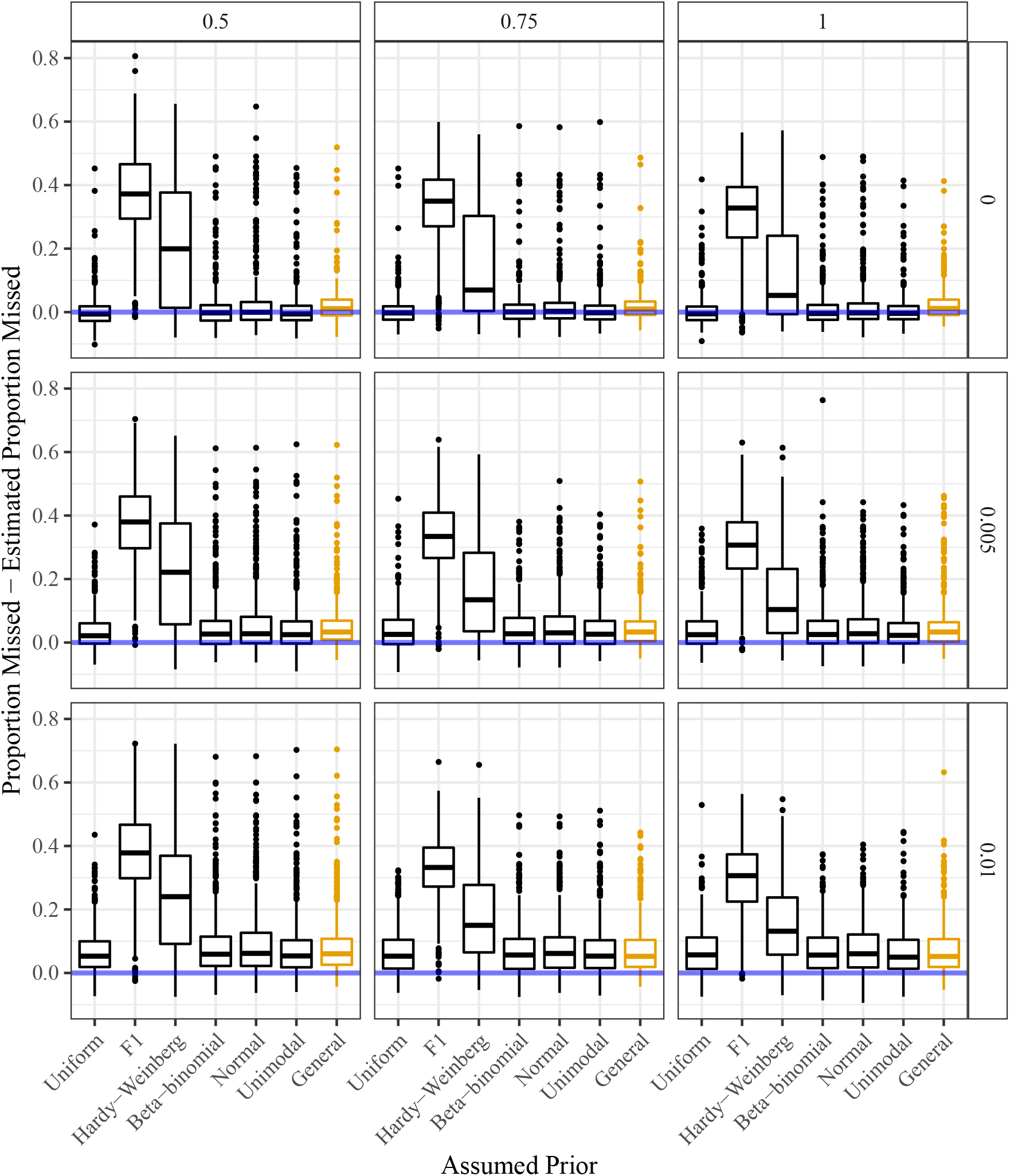
Estimated proportion of individuals incorrectly genotyped subtracted from the actual proportion of individuals incorrectly genotyped (*y*-axis) stratified by the assumed class of prior distributions (*x*-axis). The horizontal line (at *y* = 0) indicates unbiased estimation of the misclassification error rate. Column facets index varying levels of allele bias (with 1 indicating no bias) and row facets index varying levels of overdispersion (with 0 indicating no overdispersion). The genotypes were generated from a distribution designed to be favorable to the class of general prior distributions (orange).

**Figure S21:**
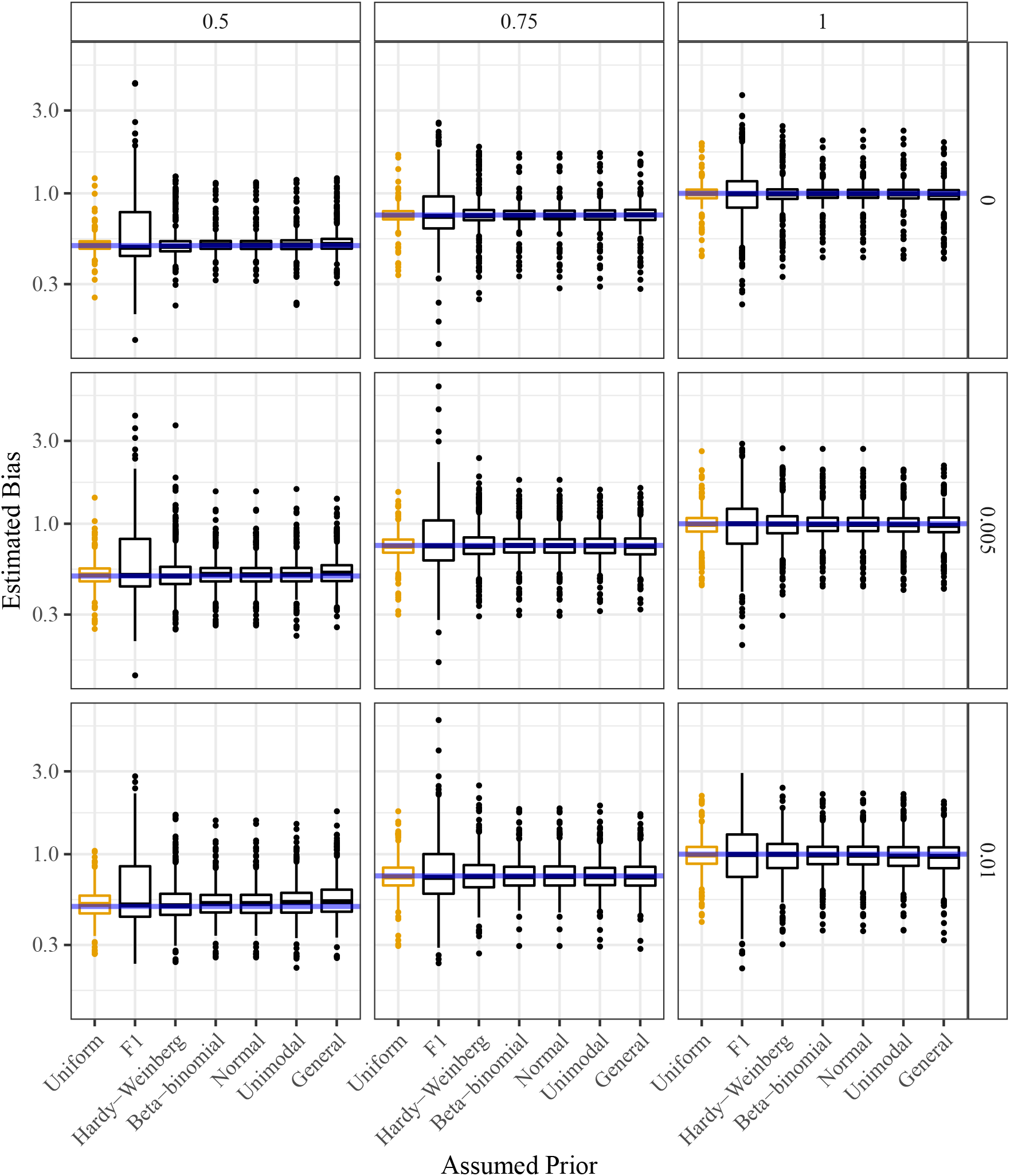
Estimated allele bias (*y*-axis) stratified by the assumed class of prior distributions (*x*-axis). The horizontal line is the actual allele bias. Column facets index varying levels of allele bias (with 1 indicating no bias) and row facets index varying levels of overdispersion (with 0 indicating no overdispersion). The genotypes were generated from a distribution designed to be favorable to the discrete uniform prior distribution (orange).

**Figure S22:**
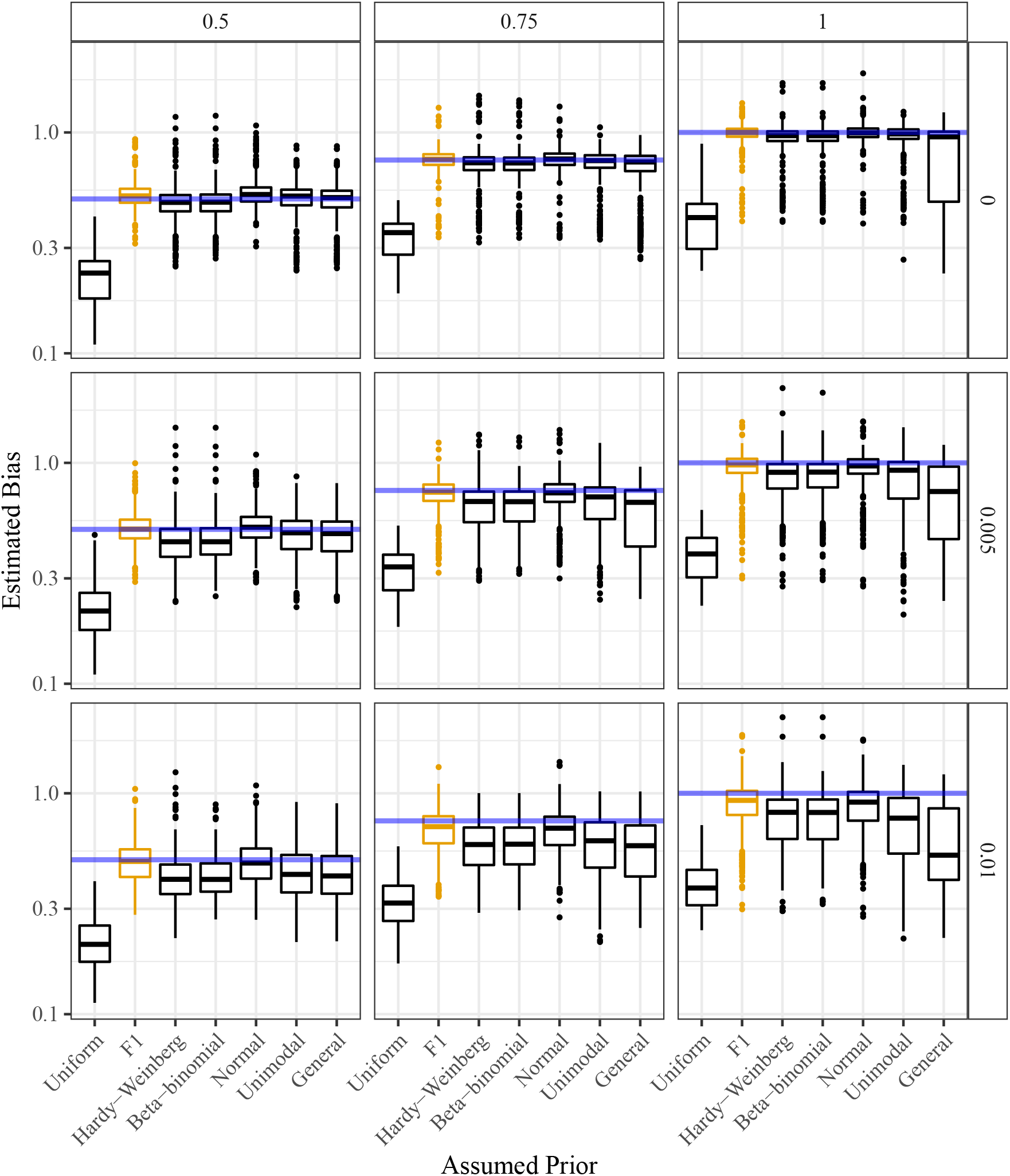
Estimated allele bias (*y*-axis) stratified by the assumed class of prior distributions (*x*-axis). The horizontal line is the actual allele bias. Column facets index varying levels of allele bias (with 1 indicating no bias) and row facets index varying levels of overdispersion (with 0 indicating no overdispersion). The genotypes were generated from a distribution designed to be favorable to the class of F1 prior distributions (orange).

**Figure S23:**
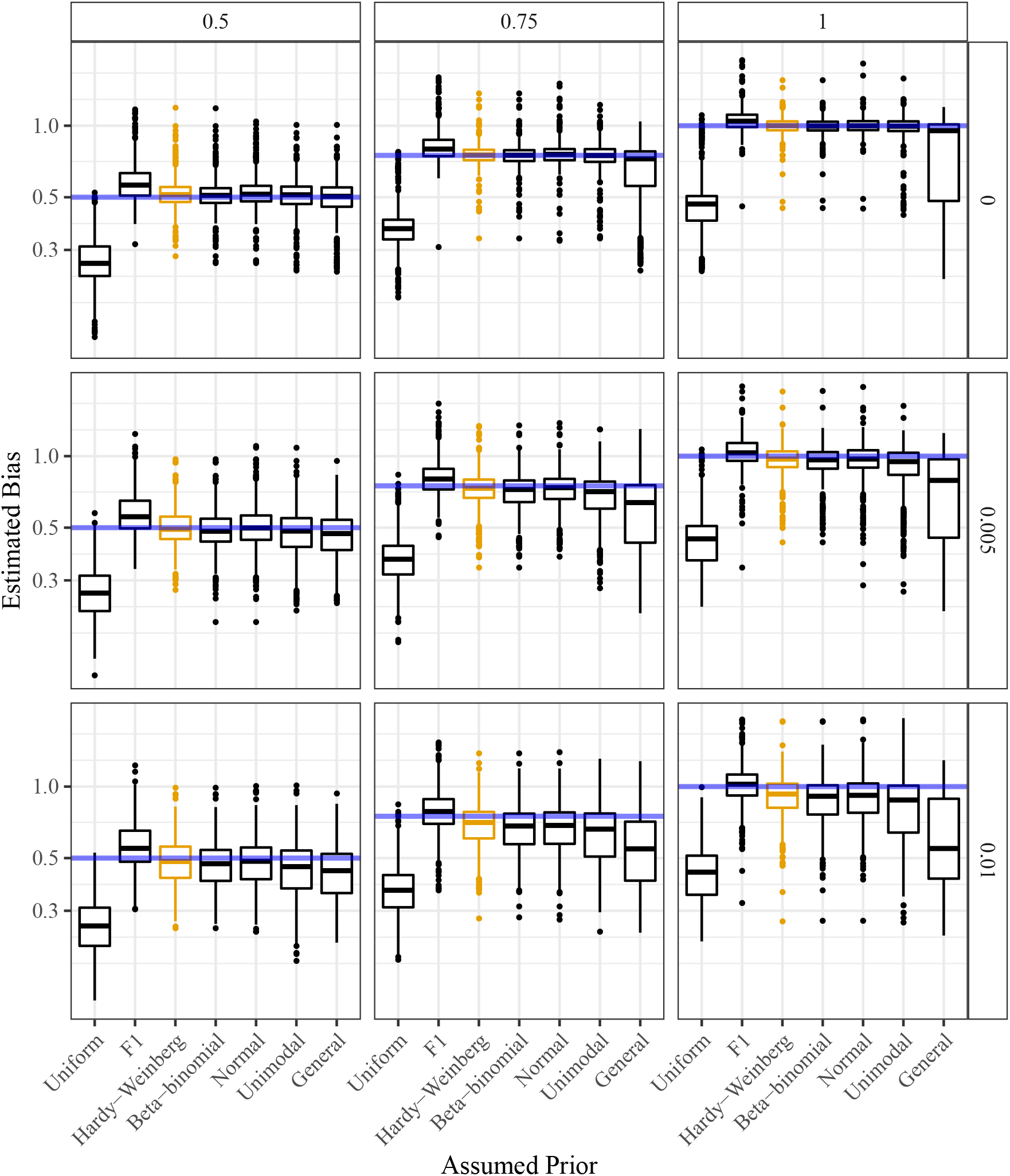
Estimated allele bias (*y*-axis) stratified by the assumed class of prior distributions (*x*-axis). The horizontal line is the actual allele bias. Column facets index varying levels of allele bias (with 1 indicating no bias) and row facets index varying levels of overdispersion (with 0 indicating no overdispersion). The genotypes were generated from a distribution designed to be favorable to the class of binomial (Hardy-Weinberg) prior distributions (orange).

**Figure S24:**
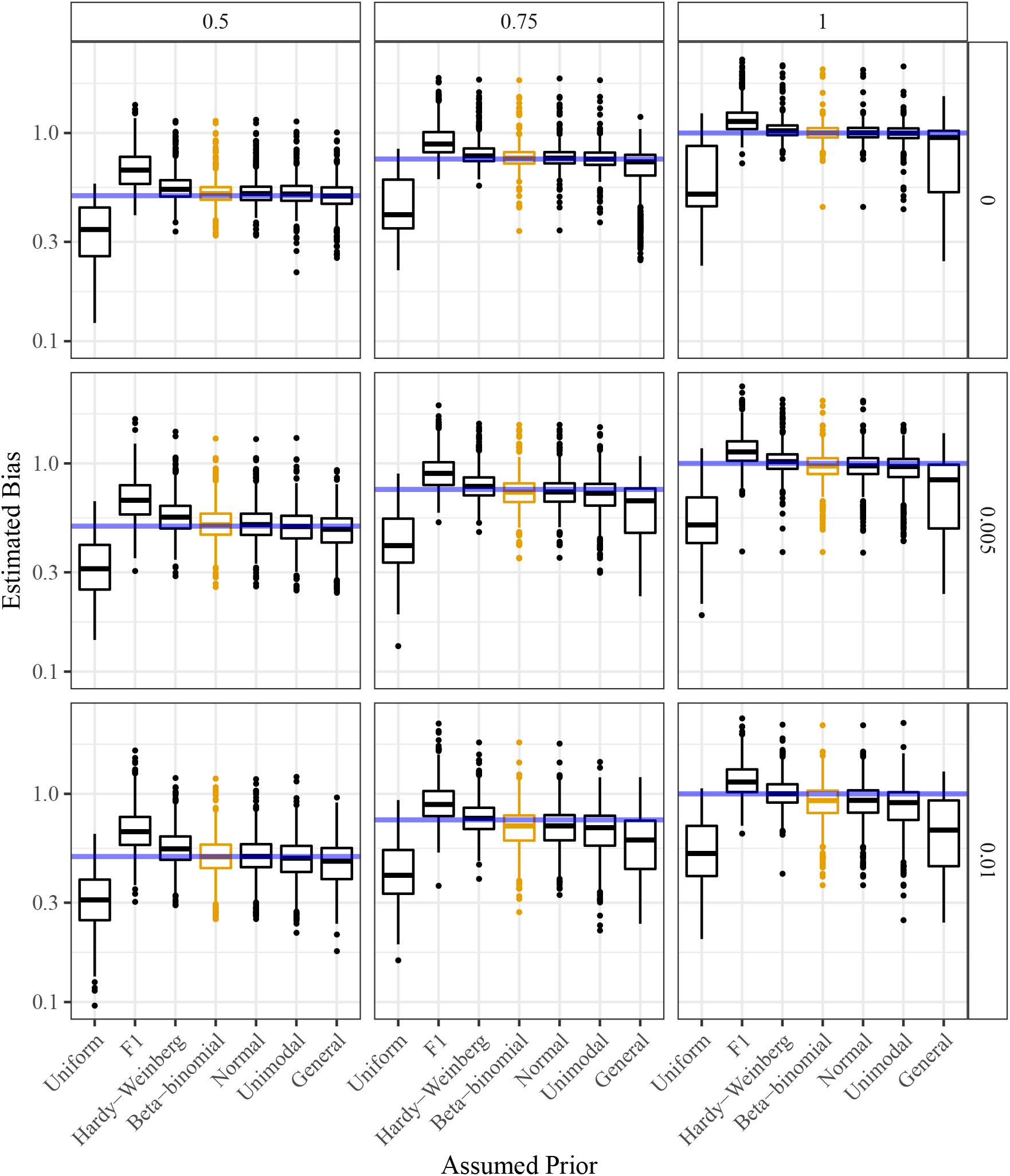
Estimated allele bias (*y*-axis) stratified by the assumed class of prior distributions (*x*-axis). The horizontal line is the actual allele bias. Column facets index varying levels of allele bias (with 1 indicating no bias) and row facets index varying levels of overdispersion (with 0 indicating no overdispersion). The genotypes were generated from a distribution designed to be favorable to the class of beta-binomial prior distributions (orange).

**Figure S25:**
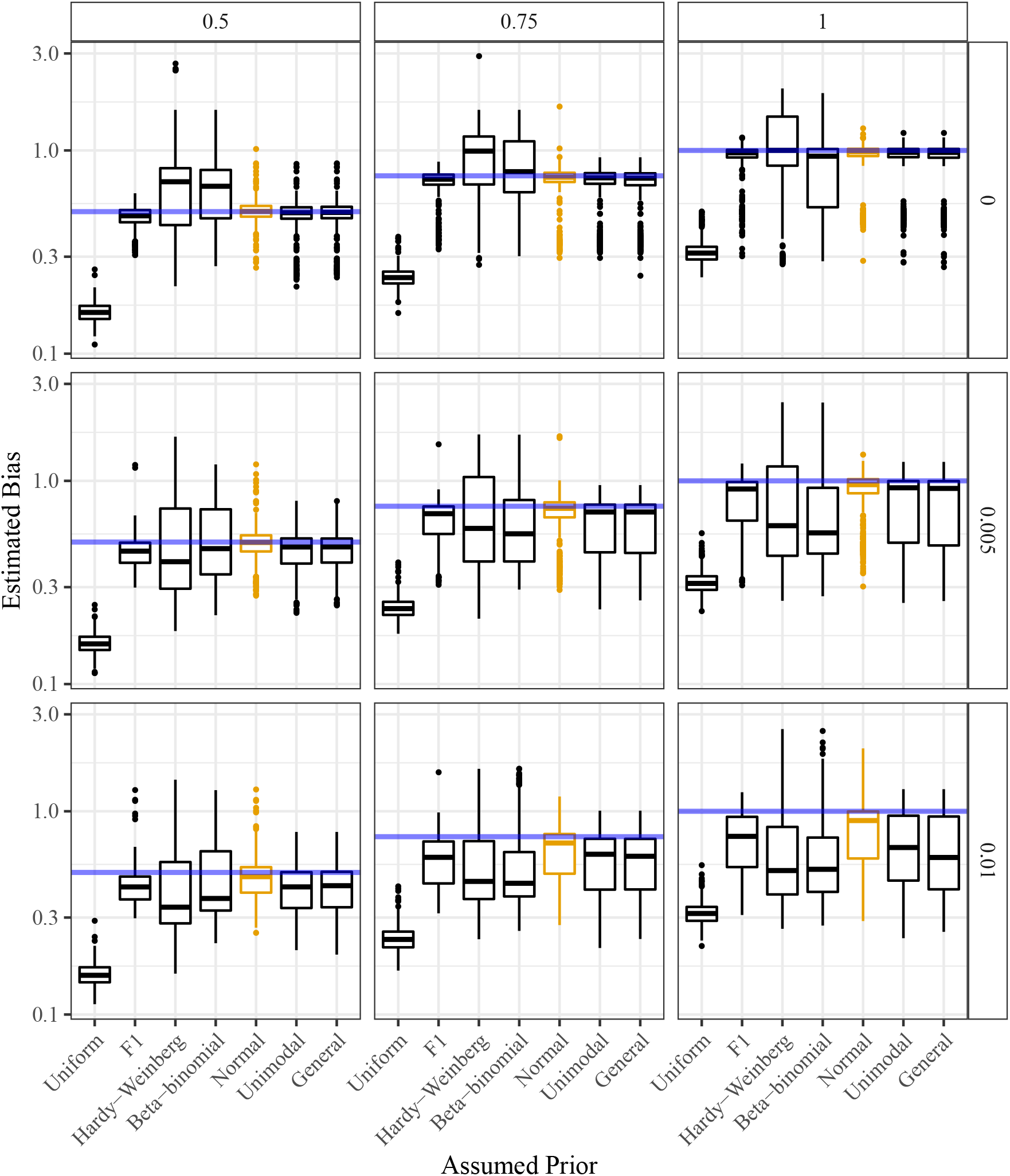
Estimated allele bias (*y*-axis) stratified by the assumed class of prior distributions (*x*-axis). The horizontal line is the actual allele bias. Column facets index varying levels of allele bias (with 1 indicating no bias) and row facets index varying levels of overdispersion (with 0 indicating no overdispersion). The genotypes were generated from a distribution designed to be favorable to the class of proportional normal prior distributions (orange).

**Figure S26:**
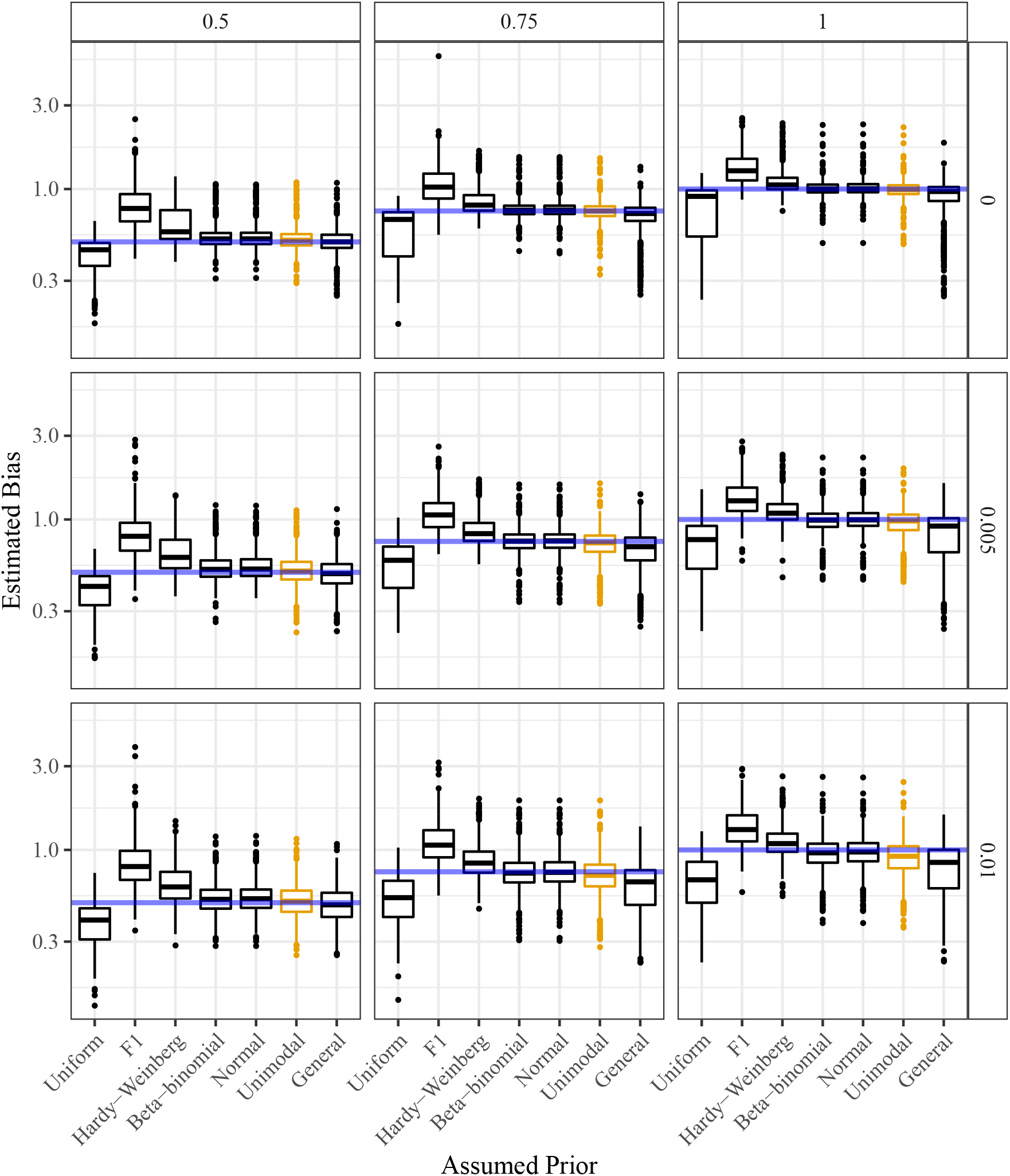
Estimated allele bias (*y*-axis) stratified by the assumed class of prior distributions (*x*-axis). The horizontal line is the actual allele bias. Column facets index varying levels of allele bias (with 1 indicating no bias) and row facets index varying levels of overdispersion (with 0 indicating no overdispersion). The genotypes were generated from a distribution designed to be favorable to the class of unimodal prior distributions (orange).

**Figure S27:**
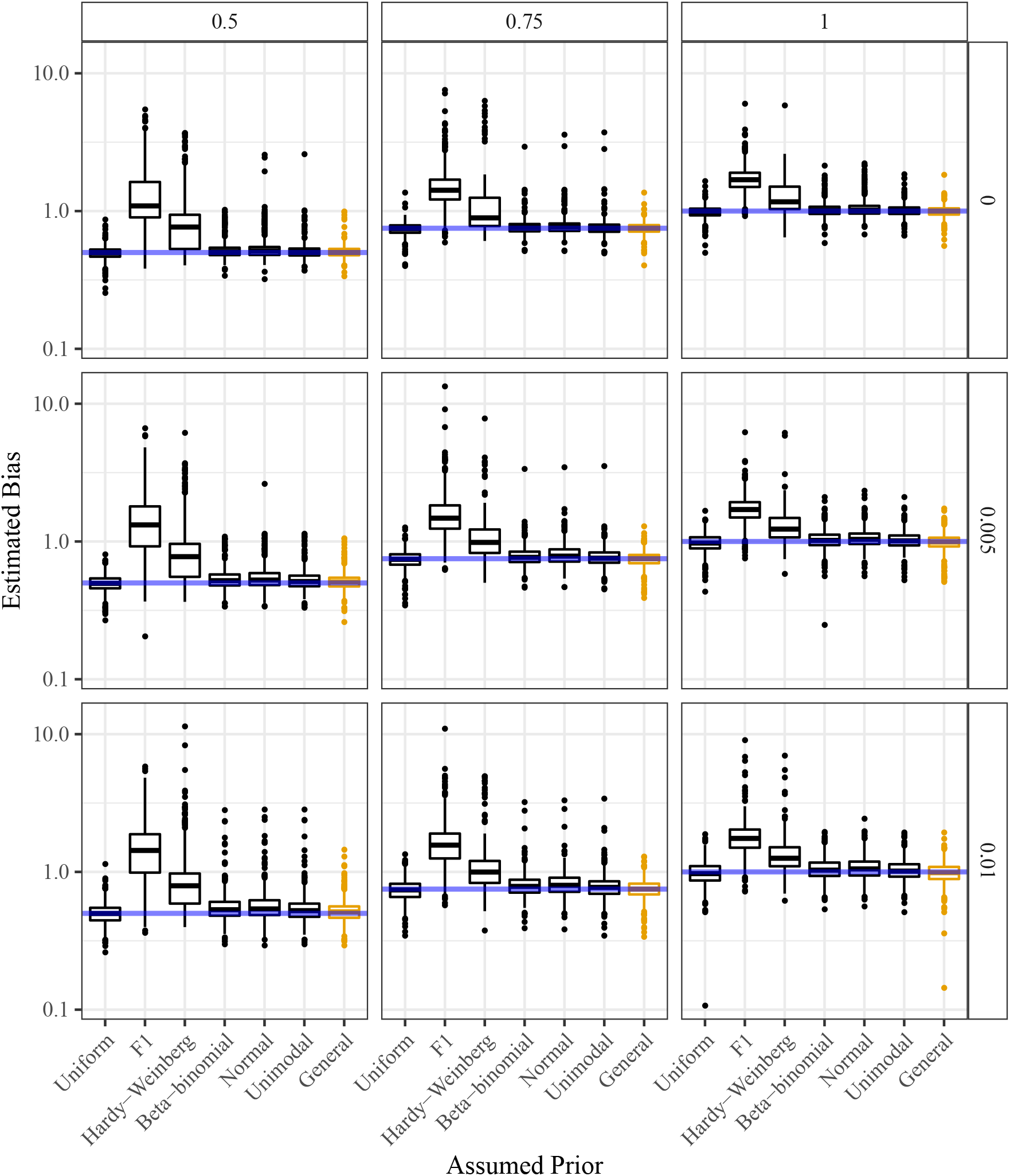
Estimated allele bias (*y*-axis) stratified by the assumed class of prior distributions (*x*-axis). The horizontal line is the actual allele bias. Column facets index varying levels of allele bias (with 1 indicating no bias) and row facets index varying levels of overdispersion (with 0 indicating no overdispersion). The genotypes were generated from a distribution designed to be favorable to the class of general prior distributions (orange).

**Figure S28:**
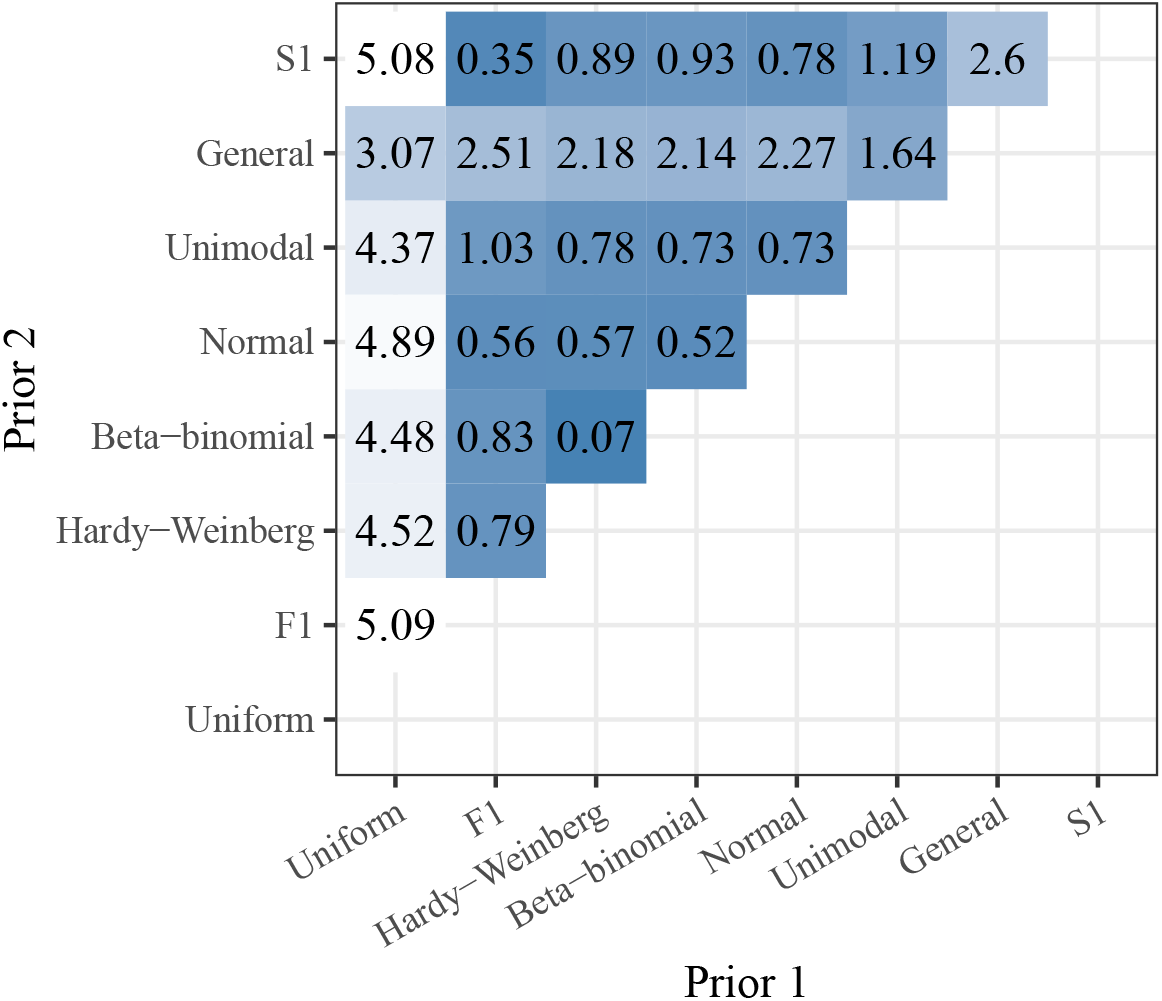
Mean Euclidean distance between posterior mean genotypes (color) resulting from two classes of prior distributions (*x* and *y* axes). The “S1” prior is the true genotype distribution, so methods perform better when their posterior mean genotypes are closer to the S1 posterior mean genotypes (mean Euclidean distance is closer to 0).

**Figure S29:**
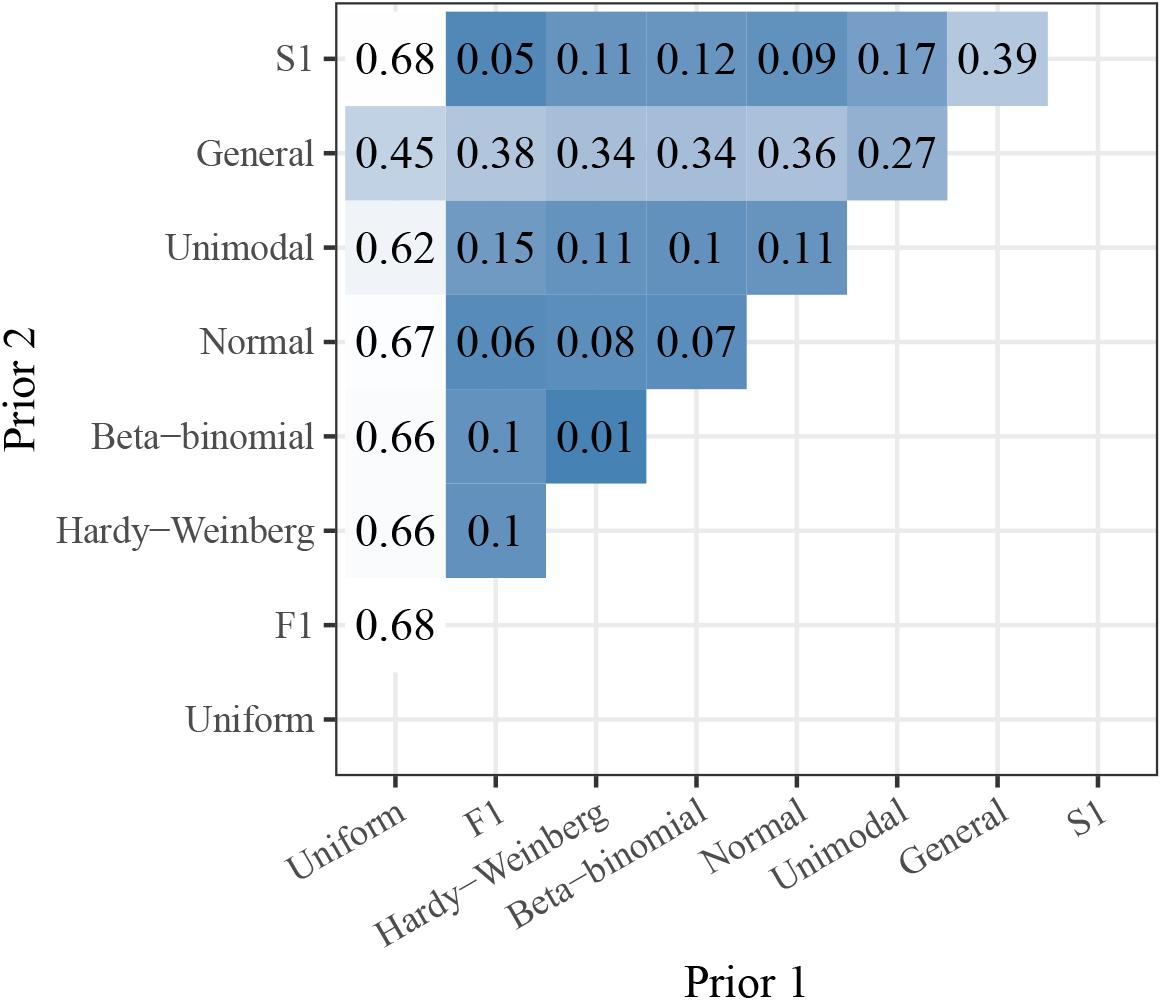
Mean proportion of posterior mode genotypes that are different (color). These posterior mode genotypes result from one of two classes of prior distributions (*x* and *y* axes). The “S1” prior is the true genotype distribution, so methods perform better when their genotype estimates are closer to those resulting from an S1 prior (the mean proportions are closer to 0).

